# Dynamic organization of visual cortical networks inferred from massive spiking datasets

**DOI:** 10.1101/2023.08.08.552512

**Authors:** Colin Graber, Yurii Vlasov, Alexander Schwing

**Affiliations:** University of Illinois Urbana Champaign, Department of Electrical and Computer Engineering

## Abstract

Complex cognitive functions in a mammalian brain are distributed across many anatomically and functionally distinct areas and rely on highly dynamic routing of neural activity across the network. While modern electrophysiology methods enable recording of spiking activity from increasingly large neuronal populations at a cellular level, development of probabilistic methods to extract these dynamic inter-area interactions is lagging. Here, we introduce an unsupervised machine learning model that infers dynamic connectivity across the recorded neuronal population from a synchrony of their spiking activity. As opposed to traditional population decoding models that reveal dynamics of the whole population, the model produces cellular-level cell-type specific dynamic functional interactions that are otherwise omitted from analysis. The model is evaluated on ground truth synthetic data and compared to alternative methods to ensure quality and quantification of model predictions. Our strategy incorporates two sequential stages – extraction of static connectivity structure of the network followed by inference of temporal changes of the connection strength. This two-stage architecture enables detailed statistical criteria to be developed to evaluate confidence of the model predictions in comparison with traditional descriptive statistical methods. We applied the model to analyze large-scale in-vivo recordings of spiking activity across mammalian visual cortices. The model enables the discovery of cellular-level dynamic connectivity patterns in local and long-range circuits across the whole visual cortex with temporally varying strength of feedforward and feedback drives during sensory stimulation. Our approach provides a conceptual link between slow brain-wide network dynamics studied with neuroimaging and fast cellular-level dynamics enabled by modern electrophysiology that may help to uncover often overlooked dimensions of the brain code.

## INTRODUCTION

Information processing in a mammalian brain is distributed across many anatomically and functionally distinct areas^1^. The inter-areal functional and anatomical connectivity can be reconstructed using electron microscopy^2^, gene co-expression^3^, projection tracing methods^4,5^, and functional neuro-imaging^6^. Further analysis of the temporal variation of neural activity across this connectome^7^ using functional neuroimaging^8–11^ have indicated that complex cognitive functions rely on highly dynamic routing of neural activity within this anatomically constrained structural backbone^12^. Neuroimaging, however, captures activity averaged over large populations (over 500K neurons per voxel) at relatively slow time scales (minutes to hours) that typically shows relatively low dimensionality^13^ hence hindering mechanistic explanation. Further insights into dynamic routing of information flows across the brain are expected with adoption of recent technological advances in multielectrode arrays^14,15^ and volumetric 2-photon imaging^16^ that enable simultaneous recording from large neuronal populations at a cellular level with single spike temporal resolution that reveals rich and high-dimensional dynamic interactions^17,18^. Descriptive statistical approaches^15,19^ that summarize the pairwise millisecond-timescale synchrony of spiking within recorded neuronal population indicate a directional flow of functional connectivity^15^ ascending along the anatomical hierarchy^5^ and reveal distinct engaged modules across the cortical hierarchy^19–21^. However, both descriptive statistical^15,19^ as well as modern probabilistic models^22–24^ typically extract interactions as time-independent “static” variables^15,19,25^ or consider temporal dynamics of the whole population^20,21,26–29^. Temporal changes in the magnitude of pair-wise correlations calculated with descriptive statistical methods have been used to track the dynamics of synaptic interactions during behavior and formation of place fields in hippocampal networks^30–32^. Temporal fluctuations of the synaptic efficacy were also inferred from pair-wise correlations in spike trains using various extended generalized linear models (GLM)^33–37^. However, the synaptic efficacy is typically calculated separately for each neuron pair while in brain networks each neuron is connected to thousands of others.

Here, to uncover the dynamic routing of information in cortical networks, we develop an unsupervised probabilistic model - Dynamic Network Connectivity Predictor (DyNetCP). As opposed to previous approaches that focus on extracting pair-wise correlations^30–37^, we aim at learning dynamic changes in connection strength for all recorded neurons in the network. Our approach is also fundamentally different from most of latent variable models that infer the dynamics of the entire recorded population in a low-dimensional latent feature space^26–29^. Instead, to enable interpretation as directed time-dependent network connectivity across distinct cellular-level neuronal ensembles, DyNetCP extracts latent variables that are assigned to individual neurons.

## MODEL

### Model architecture

Classical machine learning approaches like variational autoencoders^26,38^ successfully train dynamic models of brain activity across entire recorded neural population. However, development of such dynamic models to uncover cellular-level dynamic connectivity structure, is facing a major problem of the virtual absence of ground-truth (GT) data either as verified experimental or synthetic datasets with known time-varying connectivity. In this situation optimization of the model hyper-parameters, control of overfitting, and model verification is largely becoming a surmise. Therefore, to address this challenge, here, learning of the directed dynamic cellular-level connectivity structure in a network of spiking neurons is divided into two stages (Fig.1A). First stage, the static component, infers the static connectivity weight matrix **W**_**st**_ (Fig.1A, left) via a generalized linear model (GLM). At this stage, the classical descriptive statistical approaches like pair-wise jitter-corrected cross-correlograms^15,30^ can be used as a GT to verify the model performance. Next, in the second stage, **W**_**st**_ is used to influence the dynamic offset weight term **W**_**off**_ learned via a recurrent long-short-term-memory (LSTM) algorithm (Fig.1A, right). The final dynamic weight **W**_**dyn**_ = **W**_**st**_ + **W**_**off**_ is averaged across all trials. At this stage, the model **W**_**dyn**_ can be verified against another classical descriptive statistical method - pairwise Joint Peristimulus Time Histogram^39^ (JPSTH) that represents the time variation of joint correlations. Therefore, dividing the problem into two stages enables independent verification of static weights and dynamic offsets against GT synthetic and experimental data as well as a direct comparison of DyNetCP performance to existing models.

**Figure 1.**
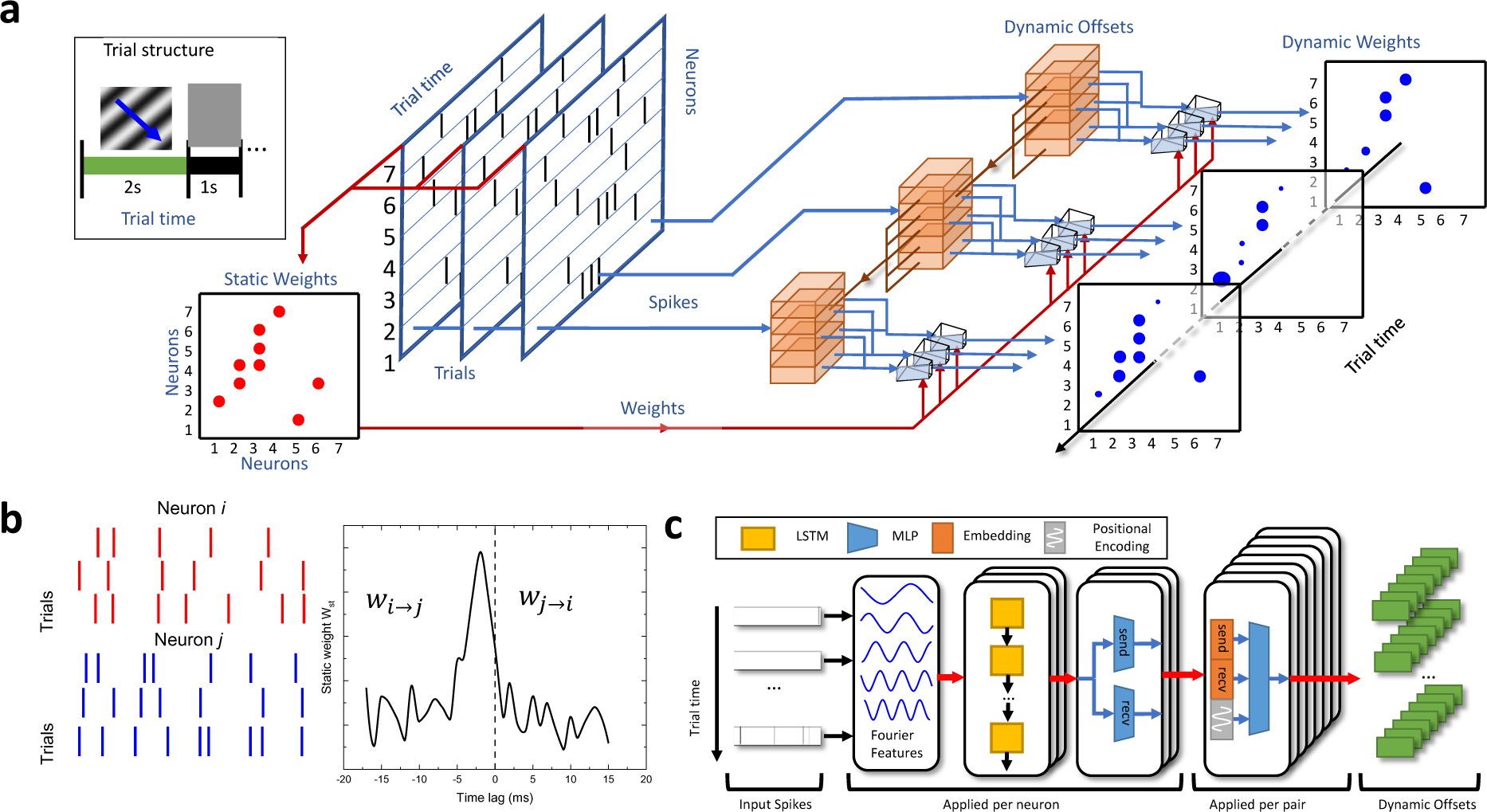
Model architecture and components design. **A)** Two-stage model architecture consisting of a static and a dynamic part. **Inset:** structure of the “drifting grating” visual stimulation trial. **B)** Schematics of spike trains recorded from neuron *i* and neuron *j* for three trials and corresponding output of the static part of the model **W**_**st**_. **C)** Schematics of the dynamic part of the mode consisting of a Fourier features transformation, followed by recurrent long-short-term-memory model, multilayer perceptron, embedding, and positional encoding.

**Fig.1 - figure supplement 1.**
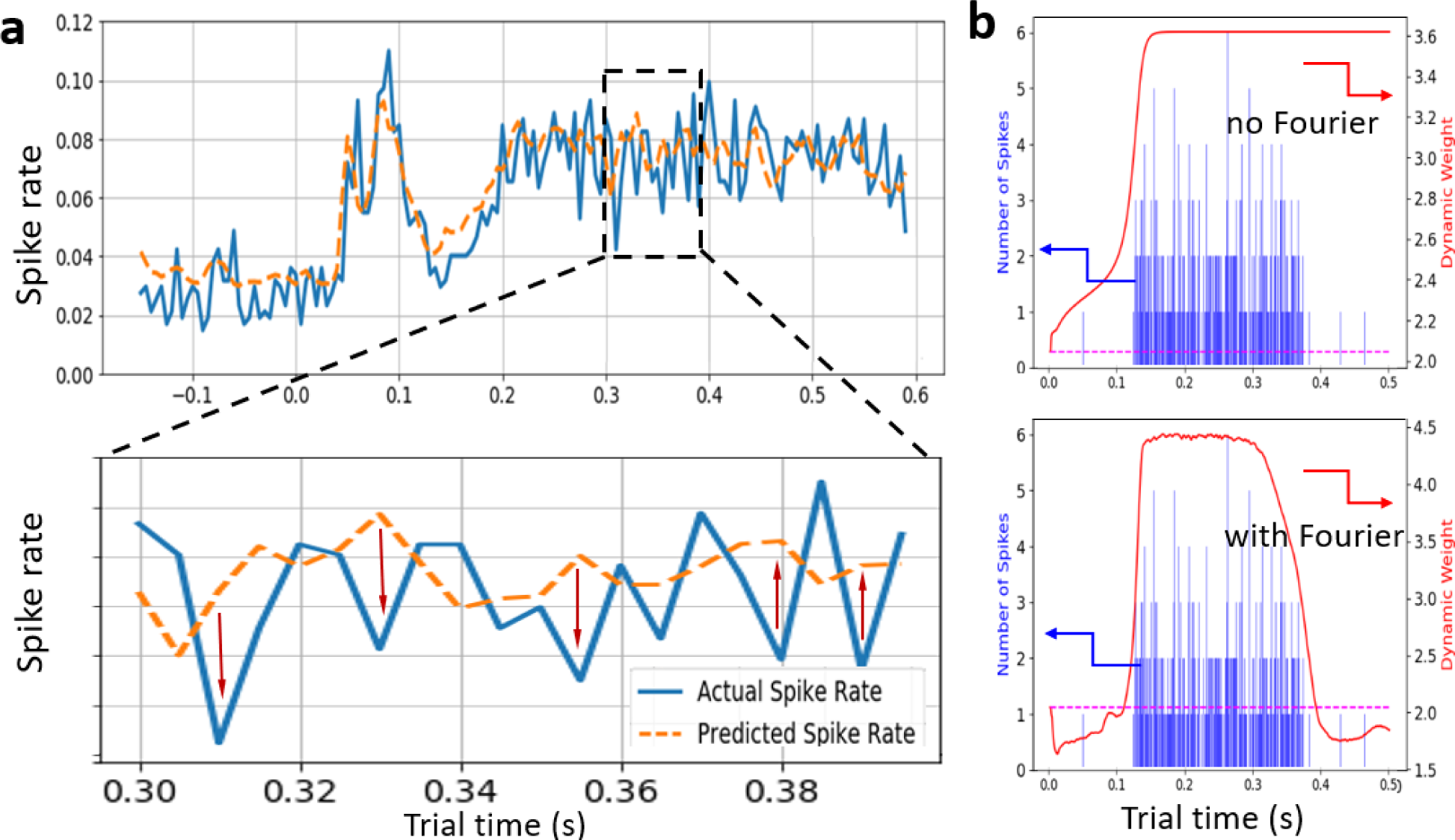
Model training and implementation. **A) Loss optimization**. Blue curve is actual spike rate for a representative unit taken from the VCN dataset (see below) as a function of trial time. The dotted yellow curve is a spike rate predicted by DyNetCP model during training. **Inset:** Loss function is designed to push the predicted spike rate to approach the actual spike rate during training. **B) Model performance with and without Fourier features.** Blue bars (left y-axis) represent a histogram of joint spiking probability (JPSTH anti-diagonal) for an example neuron pair with direct connection taken from the synthetic dataset (see below). Red curves (right y-axis) correspond to trial-averaged learned dynamic weights, while dotted magenta line represents static weight value. DyNetCP model without Fourier features (top plot) failed to properly localize the period where induced spiking is occurring, while use of Fourier features (bottom plot) enables to do so correctly.

The model operates on trial-aligned binned neural spikes. We collect *N* neurons recorded for *T* time bins binned at *d* bin width across *M* trials in tensor *X* ∈ {0,1}^*M*×*N*×*T*^, where each entry 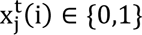 represents the presence or absence of a spike from neuron *j* at time *t* in trial *i*. The spiking probability 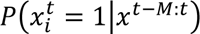 of neuron *i* at time *t* is modelled as follows:

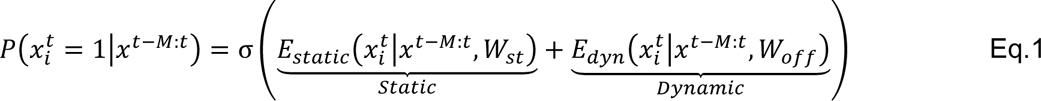

where σ is the sigmoid function, *E*_*static*_ ∈ ℝ and *E*_*dyn*_ ∈ ℝ are static and dynamic connectivity logits depending on the trainable weight matrix **W**_**st**_ and the dynamic offset model **W**_**off**_ respectively. Given spiking data as input, the model produces two sets of output weight matrices. The first set, **W**_**st**_, encodes an estimate of the static network structure and contains one weight per pair of neurons averaged across the trial time. These represent the time-averaged influence the spikes of the whole population have on spiking probability of a given neuron. The second set, **W**_**dyn**_, contains a single weight for every pair of neurons for every point in time during the experimental trial. These weights, which are a function of the input spiking data of all neurons in a population, represent a time-varying offset to the static weights. Hence, they encode temporal changes in activity patterns, and are used to determine when a given connection is being active and when it is unused. Therefore, in this formulation, the model learns time-varying connectivity patterns that can be directly linked to the underlying network structure at a cellular level.

### Static connectivity inference

The static component of DyNetCP (Fig.1A, left), i.e.,

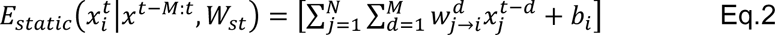

learns static connectivity weights 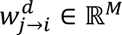 that encode static network structure averaged over the time of the trial *t*, and *b*_*i*_ ∈ ℝ are biases. We adopt a multivariate autoregressive GLM model^23^ without temporal kernel smoothing that learns **W**_**st**_ as a function of a time lag *d* (Fig.1B). Sharp peak at short time lags implies increased probability of spiking of neuron *j* when neuron *i* is producing a spike that can be an indication that these two neurons are synaptically connected. An apparent time lag of the low-latency peak indicates a functional delay and defines the directionality of the putative connection. The static component of DyNetCP is implemented for all neurons jointly using a 1-dimensional convolution. Each convolutional filter contains the static weights used to predict the spiking probability for a single neuron as a function of the spiking history of all other neurons. The bias term *b*_*i*_ in Eq.2 is initialized to the logit function σ (i.e., inverse sigmoid) applied to the spike rate of each neuron calculated for a given time bin.

### Dynamic connectivity inference

The dynamic component of DyNetCP (Fig.1A, right), i.e.,

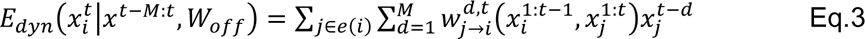

learns the dynamic connectivity weights 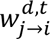 which are a function of input spike trains 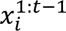 and all other 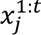. Each of these weights represents an offset to the time-averaged static weight at time *t* for a connection at time lag *d* (Eq.2). Hence, assuming the spikes of neuron *i* at a time lag *d* are influenced by spikes from all 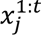, the total influence applied to the spiking of neuron *i* from neuron *j* at time *t* is 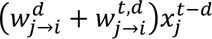. Unlike the static weights, these dynamic weights are allowed to change throughout an experimental trial as a function of the spiking history of the connected neurons.

The architecture of the dynamic part (Fig.1C) consists of several stages including generation of Fourier features, recurrent long-short-term-memory (LSTM) model, multilayer perceptron (MLP), embedding, and positional encoding. First, we shift each spike train such that each entry lies in {−1, 1} and then represent each spike train using Fourier features^40^ that were recently introduced in the computer vision domain to facilitate the learning of high-frequency functions from low-dimensional input. Specifically, we encode every spike 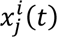 using the following transformation:

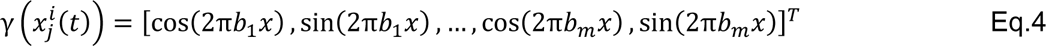

Here, we use *m* = 64 features, where coefficients *b*_1_, …, *b*_*m*_ are linearly spaced between –1 and 1. We evaluate the use of Fourier features within the model on synthetic data consisting of a simulated network of neurons, where one neuron induces spiking in a second neuron during the middle 250 ms of each trial (Fig.1 - figure supplement 1B). DyNetCP without Fourier features is unable to properly localize the period where induced spiking is occurring, while the model with Fourier features can do so correctly.

The spike train features for each neuron are then input into separate LSTMs to produce an embedding per neuron per time step. Each of these LSTMs uses a separate set of weights and has a hidden dimension of 64. The LSTM embeddings are passed through two MLPs, *f*_send_ and *f*_recv_, where *f*_send_ produces an embedding which represents the outgoing neural signal to other neurons and *f*_recv_ produces an embedding which represents the incoming signal from other neurons. Each of these MLPs consists of one linear layer with output size 64 and a ReLU activation. For each pair *n*_*j*_ → *n*_*i*_ of neurons being modeled, the send embedding ℎ_*s*,*j*_(*t*) is concatenated with the receive embedding ℎ_*r*,*i*_(*t* − 1) as well as a time encoding ℎ_*T*_(*t*) which represents the trial time and is represented as

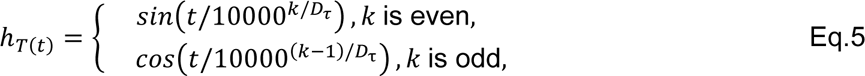

where *D*_τ_ is the size of the positional encoding which we set to 128. Note that the send embedding contains spiking information through time *t*, i.e., it contains information about the most recent spikes, which is necessary to properly represent synchronous connections. Importantly, the receive embedding only contains spiking information through time *t* − 1 such that the output spiking probability is not trivially recoverable. Finally, this concatenated embedding is passed through a 2-layer MLP with hidden size 384 and a Rectified Linear Unit (ReLU) activation, producing the dynamic weight vector 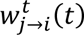 which contains dynamic weight offsets for each delay *d* ∈ {1, …, *M*}.

### Model training

The loss used to train the model consists of two terms. The first is a negative log likelihood over the predicted spike probabilities, which encourages the weights to accurately encode the influence of spiking from other neurons on the spiking of the neuron in question. The entire loss is written as

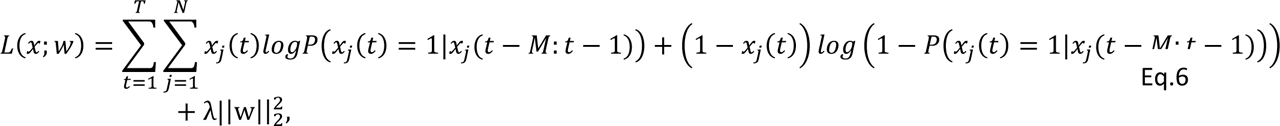

where *w* represents all the predicted static and dynamic weights, and λ is a coefficient used to balance the magnitudes of the losses. Loss optimization during training results in predicted spike rate (PSTH) that is approaching the actual spike rate (Fig.1 – figure supplement 1A).

We introduce a squared *L2* penalty on all the produced static and dynamic weights as the second term to discourage the network from overpredicting activity for times when no spikes are present 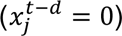 and the weight multiplied by zero is unconstrained. In these cases, adding the penalty provides a training target. For all experiments we used an *L2* weight between 0.1 and 1 that produced the best fit to the descriptive statistics results discussed in detail below (Methods).

Overall, once the architecture of Fig.1C is established, there are just a few parameters to optimize including L*2* regularization and loss λ coefficient. Other variables, common for all statistical approaches, include bin sizes in the lag time and in the trial time. Decreasing the bin size will improve time resolution while decreasing the number of spikes in each bin for reliable inference. Therefore, the number of spikes threshold and other related thresholds need to be adjusted as discussed in detail in the Methods, Section 4.

## RESULTS

The DyNetCP model is designed to be trained in a single step to produce both static and dynamic weight matrices simultaneously for the whole dataset. However, for the purpose of this work, we are interested in verification and detailed comparative analysis of the model performance that enables clear justification for the choice of model parameters. Therefore, here, training of the model to analyze a new dataset is implemented in two separate stages. In the first stage, only the static weights **W**_**st**_ are trained for all the valid neurons while the dynamic weights **W**_**dyn**_ are all set to zero. In the second stage, the static model parameters and static weights are fixed, and we train only the dynamic component of the model.

This two-stage architecture enables detailed comparison of static and dynamic matrices to classical descriptive statistical methods as well as to modern probabilistic models. In the virtual absence of synthetic or experimental datasets with known GT such comparisons are becoming crucial for tuning the model parameters and for validation of the results. It is especially important for validation of the dynamic stage of the DyNetCP since it predicts **W**_**dyn**_ that describes temporal variation of the connection strength that is assigned to individual neurons, and, therefore, could not be directly compared to most of the population dynamics models that produce temporal trajectories in a latent feature space of the whole population^26–29^. In what follows we consider these comparisons separately.

### Static DyNetCP can faithfully reproduce descriptive statistics cross-correlogram

The DyNetCP formulation of its static stage (Eq.2) is similar to classical cross-correlogram^15,30^ (CCG) between a pair of neurons *n*_1_ and *n*_2_ that is defined as:

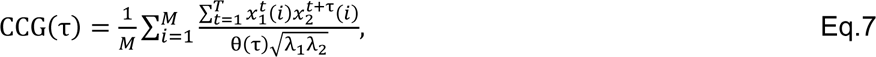

where τ represents the time lag between spikes from *n*_1_ and *n*_2_, λ_*j*_ is the spike rate of neuron *j*, and θ(τ) is a triangular function which corrects for the varying number of overlapping bins per delay. However, instead of constructing the statistical CCG histogram for each pair individually, the model given in Eq.2 infers coupling terms from spiking history of all *N* neurons jointly. Typically, to minimize the influence of a potential stimulus-locked joint variation in population spiking that often occurs due to common input from many shared presynaptic neurons, a jitter correction^15,30,41–43^ is applied. To directly compare the results of static DyNetCP with jitter-corrected CCG, we train two static models (Methods) – one on original spike trains and the second on a jitter-corrected version of the data generated using a pattern jitter algorithm^41^ with a jitter window of 25ms.

For this comparison we used a subset of the Allen Brain Observatory-Visual Coding Neuropixels (VCN) database^15^ that contains spiking activity collected in-vivo from 6 different visual cortical areas in 26 mice (Methods, Table 1) passively viewing a “drifting grating” video (top left inset in Fig.1A). For a pair of local units (spatially close units from Layer 5 of the VISam area) comparison of static DyNetCP (Fig.2A, red curve) with jitter-corrected CCG (Fig.2A, black curve) shows close correspondence (Pearson correlation coefficient r_p_=0.99 and p-value P_p_<0.0001). Residual analysis (Fig.2A, bottom panel) shows a mean value of 0.01±0.02 SEM (Fig.2A, top right panel) that is smaller than the CCG values on the flanks (bottom right panel) (mean:0.11±0.06 SEM). For such a local connection the peak is narrow (∼2ms) with a functional delay of −2.3ms. For a pair of spatially distant units (units in VISam and VISp areas) a comparison of static DyNetCP (Fig.2B, red curve) with jitter-corrected CCG (Fig.2B, black curve) also shows close correspondence (r_p_=0.99 and P_p_<0.0001). For such a distant connection the peak is wider (∼9ms) with a functional delay of −5.8ms. Further examples of comparisons of both models for all the units in the session showed similar results (see accompanying Jupyter Notebook).

**Figure 2.**
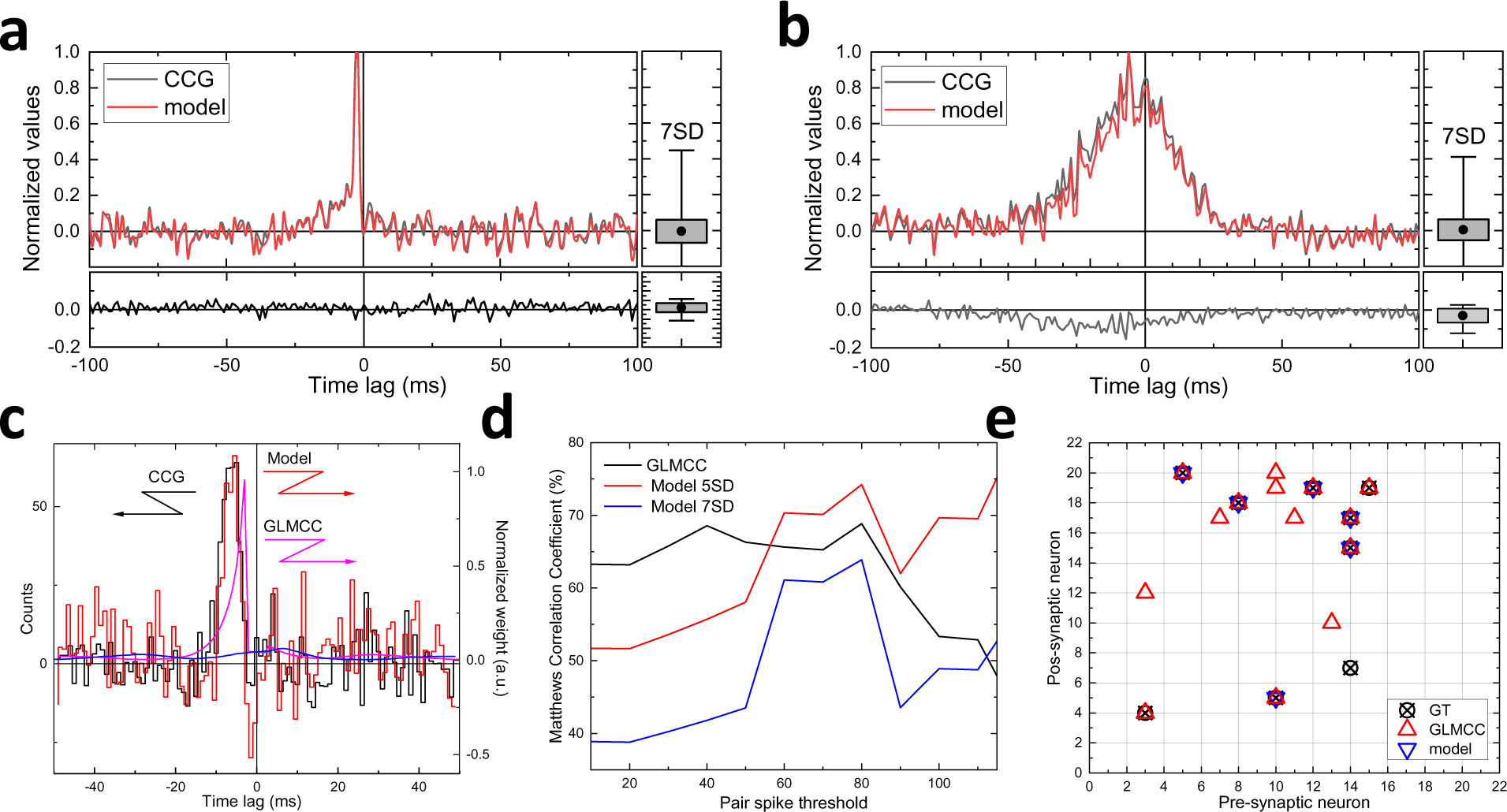
Comparison of inferred static connectivity to alternative methods. **A)** Comparison of static DyNetCP (red curve) with jitter-corrected CCG (black curve) for a representative pair of local units (session 771160300, spatially close units from Layer 5 of the VISam area). Residuals (bottom panel) and residual analysis (right panels). **B)** Comparison of static DyNetCP (red curve) with jitter-corrected CCG (black curve) for a representative pair of spatially distant units (session 771160300, units from VISam and VISp areas). **C)** Comparison of jitter-corrected CCG (black), GLMCC (magenta), and DyNetCP (red) weights for a pair of excitatory HH neurons from a synthetic dataset. **D)** MCC calculated for GLMCC (black) and for DyNetCP with *α*_*w*_ =5SD (red) and 7SD (blue). **E)** Static connectivity **W**_**st**_ for a subset of excitatory HH neurons with GT (crossed circles) compared to connections recovered by GLMCC (red triangles) and DyNetCP (blue triangles).

**Fig.2 - figure supplement 1.**
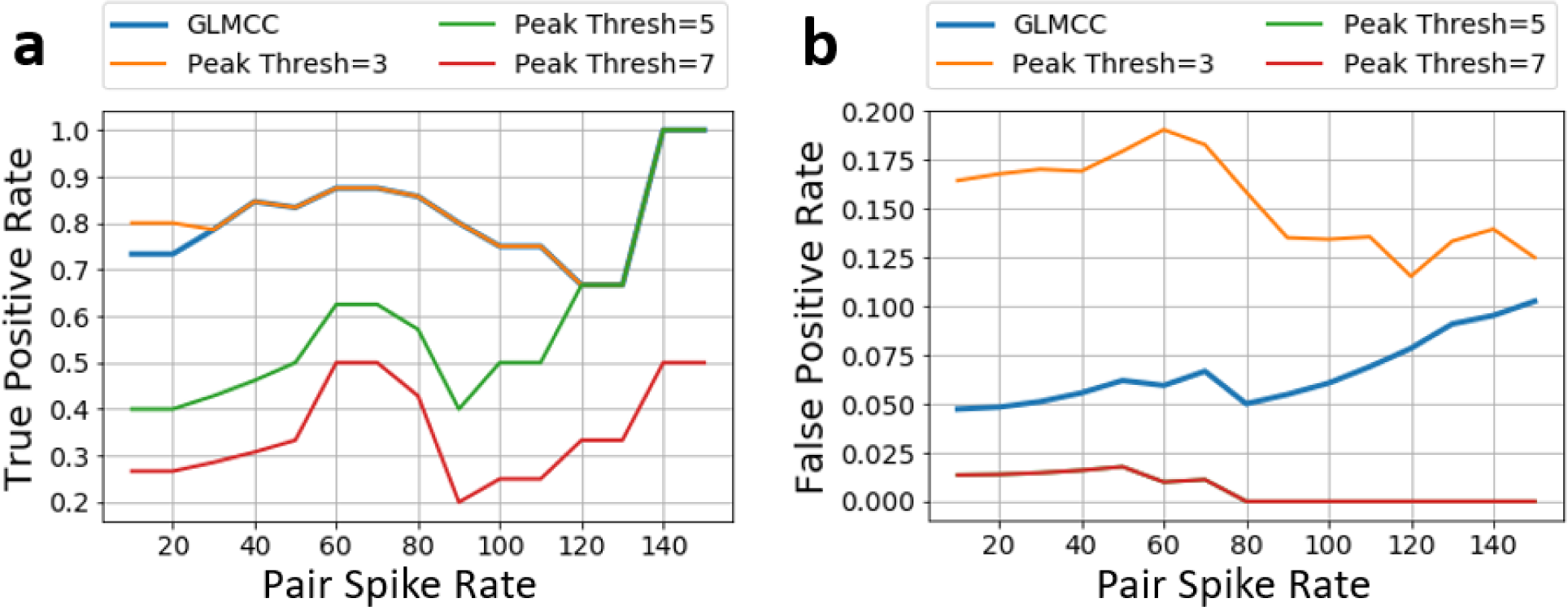
TP and FP rates as a function of pair spike threshold. **A) Recovery of True Positive (TP) connections in HH synthetic dataset**. TP connections recovered by DyNetCP model at different pair spike thresholds *α*_*p*_. Yellow, green, and red curves correspond to peak threshold taken as 3SD, 5SD, and 7SD, respectively. TP connections recovered by GLMCC model are shown by blue curve. Since a predicted connection is counted as a *TP* only once (i.e., *i* → *j* and j→ *i* do not count as two negatives if *i* is not connected to *j*) and the number of *GT* connections decreases when α_*p*_ increases, the MCC exhibits an increase beyond α_*p*_=100. **B) Generation of False Positive (TP) connections in HH synthetic dataset.** FP connections generated by the DyNetCP model at different pair spike rates. Yellow, green, and red curves correspond to peak threshold *α*_*p*_ taken as 3SD, 5SD, and 7SD, respectively. FP connections recovered by GLMCC model are shown by blue curve.

**Table. 1.** Experimental dataset recorded in-vivo in 6 visual cortices. Sessions used for model training and analysis taken from the publicly available Allen Brain Observatory-Visual Coding Neuropixels (VCN) database. We use a “functional connectivity” subset of these recordings corresponding to the “drifting grating 5 repeats” setting recorded for animals. Filtering of recorded units (see statistical analysis criteria below) and classification of valid pairs results in a significant reduction of units (5.6% of raw units or 23% of clean units) used for dynamic connectivity inference. The last two columns represent the number and percentage of trials when the animal was running with average speed over 5cm/sec

| Session | Units | Clean units | Valid Units | Valid Units % | Valid Pairs | Run | Run % |
| --- | --- | --- | --- | --- | --- | --- | --- |
| 766640955 | 842 | 145 | 25 | 17% | 24 | 50 | 10% |
| 767871931 | 713 | 185 | 53 | 29% | 49 | 146 | 30% |
| 768515987 | 802 | 198 | 41 | 21% | 43 | 4 | 1% |
| 771160300 | 930 | 223 | 35 | 16% | 35 | 115 | 24% |
| 771990200 | 546 | 108 | 42 | 39% | 41 | 43 | 9% |
| 774875821 | 649 | 172 | 43 | 25% | 12 | 126 | 26% |
| 778240327 | 784 | 275 | 78 | 28% | 56 | 413 | 86% |
| 778998620 | 793 | 233 | 44 | 19% | 15 | 240 | 50% |
| 779839471 | 863 | 203 | 29 | 14% | 17 | 4 | 1% |
| 781842082 | 833 | 229 | 36 | 16% | 38 | 66 | 14% |
| 786091066 | 700 | 306 | 17 | 6% | 13 | 408 | 85% |
| 787025148 | 696 | 134 | 18 | 13% | 21 | 110 | 23% |
| 789848216 | 415 | 81 | 31 | 38% | 26 | 183 | 38% |
| 793224716 | 781 | 198 | 82 | 41% | 77 | 171 | 36% |
| 794812542 | 1005 | 348 | 38 | 11% | 30 | 54 | 11% |
| 816200189 | 634 | 161 | 46 | 29% | 75 | 11 | 2% |
| 819186360 | 531 | 166 | 26 | 16% | 13 | 13 | 3% |
| 819701982 | 585 | 163 | 38 | 23% | 29 | 440 | 92% |
| 821695405 | 474 | 148 | 38 | 26% | 28 | 40 | 8% |
| 829720705 | 529 | 232 | 54 | 23% | 43 | 88 | 18% |
| 831882777 | 657 | 264 | 65 | 25% | 62 | 391 | 81% |
| 835479236 | 582 | 199 | 36 | 18% | 38 | 70 | 15% |
| 839068429 | 742 | 150 | 49 | 33% | 57 | 401 | 84% |
| 839557629 | 450 | 136 | 26 | 19% | 11 | 307 | 64% |
| 840012044 | 758 | 179 | 67 | 37% | 97 | 135 | 28% |
| 847657808 | 874 | 231 | 53 | 23% | 70 | 72 | 15% |
| **Total** | 18168 | 5067 | 1110 | 23% | 1020 | 4101 | 33% |

Following the common formulation, a putative excitatory (or inhibitory) synaptic connection can be considered statistically significant for both methods when the magnitude of the short-latency peak (or trough) exceeds the peak height threshold α_w_ (lower than −α_w_) compared to the standard deviation (SD) of the noise in the flanks far from the peak^15,19,25,30^ (whisker plots on top right panels in Fig.2A,B).

### Static DyNetCP reproduces GT connections on par with alternative probabilistic method

We compare the performance of DyNetCP to a recent GLM-based cross correlation method (GLMCC)^25^ that detects connectivity by fitting a parameterized synaptic potential filter (Fig.2C, magenta). For these experiments, we use the synthetic dataset^25^ consisting of 1000 Hodgkin-Huxley (HH) neurons with known GT synaptic connections (Methods). We focus specifically on recovering ground-truth (GT) excitatory connections. The **W**_**st**_ inferred by DyNetCP for a representative pair of neurons (Fig.2C, red) reproduces well the short-latency narrow peak and the flanks of the postsynaptic potential (PSP) filter learned by GLMCC (Fig.2C, magenta). Comparing to CCG histogram (Fig.2C, black), the DyNEtCP is performing much better than GLMCC (r_p_=0.67 and P_p_<0.001), however the results are not as directly interpretable as GLMCC that outputs synaptic weight from the PSP fitting. Both models, however, can infer presence or absence of the connection by examining the height of the low latency peak.

To facilitate a performance comparison, we use the Matthews Correlation Coefficient (MCC), defined as

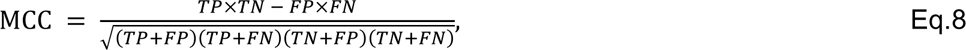

where *TP*, *TN*, *FP*, and *FN* are true positive, true negative, false positive, and false negative pair predictions, respectively. MCC optimization balances between maximizing the number of *TP* pairs recovered while minimizing the number of *FP* pairs which are predicted.

Performance of all three methods, classical CCG and probabilistic GLMCC and DyNetCP, will be strongly affected by the total number of spikes available for inference. To ensure the statistical significance of connections inferred by all 3 methods a large enough number of spikes should be present in each time bin (Methods) that is tracked by two threshold parameters: a unit average spike rate α_*s*_, and a pair spike parameter *α*_*p*_ that is the number of spikes that both units generate in a time window of interest (Methods). Fig.2D compares the performance of DyNetCP and GLMCC on a synthetic dataset, while the number of spikes present for learning is varied via a threshold pair spike parameter *α*_*p*_. Increasing *α*_*p*_ ensures statistically significant inference about the presence or absence of a connection^25^ (Methods). As α_*p*_ increases, the MCC for both models shows a maximum around α_*p*_=80 corresponding to the optimal balance. For a conservative value of *α*_*w*_=7SD (blue), the MCC for the DyNetCP model is 63%, approaching the performance of GLMCC (68%, black). When *α*_*w*_ is relaxed to 5SD (red), DyNetCP recovers a larger number of TP with a reasonable increase of FP connections (Fig.2 - figure supplement 1), increasing MCC up to 74%, thus outperforming GLMCC. Overall, as long as DyNetCP has access to a sufficient number of spikes for inference, it can restore GT static connectivity on par with or even better than GLMCC (Fig.2E).

Though GLMCC performs well in recovering static network structure and it produces directly biologically interpretable results, we emphasize that GLMCC is not compatible with modeling the network dynamics and thus cannot be integrated into DyNetCP. Specifically, our goal is to model the correlations of spiking of a network of neurons and track the changes of these correlations across time. GLMCC learns correlations as a function of the time lag relative to the spiking time of a reference neuron. It is hence unable to link correlations to specific moments in a trial time. Therefore, it cannot represent the dynamics of correlations, preventing its use in analysis of network activity relative to subject behavior. In contrast, the formulation of static DyNetCP model of Eq.2 provides a weight matrix **W**_**st**_ which is compatible with the subsequent inference of time-dependent dynamic weights **W**_**dyn**_.

### Static connectivity inferred by the DyNetCP from in-vivo recordings is biologically interpretable

Recent reconstruction of the brain-wide connectome of the adult mouse brain using viral tracing of axonal projections^4,5^ enables assignment of anatomical hierarchical score values to cortico-cortical projections across visual cortices^4^ (Fig.3A). Similarly, measuring the time lags of the maximum peaks in the CCG of recorded neuron pairs (e.g. as in Fig.1B) can be used to assess a hierarchy of functional connectivity between different cortical areas. Such analysis of functional delays extracted from massive recording of spiking activity across visual cortices^15^ revealed strong correlation of area-averaged functional delays and the hierarchical score values indicative of the hierarchical processing of visual information. Classical CCG (as well as GLMCC), however, recovers static network structure by modeling pairwise correlations and hence must be run separately for each pair in the dataset without considering the relative influence of other pairs in the network. Each neuron, however, is connected to thousands of others and is potentially contributing to multiple networks simultaneously. In contrast, DyNetCP learns connections of an entire network jointly in one pass and can track multiple connections beyond pair-wise. It is, therefore, instructional to compare the hierarchical interactions between visual cortices inferred by DyNetCP to CCG results using the same large-scale in-vivo VCN dataset^15^. Following the previous methodology^15,19^, functional delays corresponding to the time lags of jitter-corrected CCG peaks are determined for all recorded pairs across all six cortical areas with peak amplitudes above α_w_ of 7SD (5023 pairs, n=26 animals). The median time lags averaged across pairs within a given visual area (Fig.3B) are identified as the area’s functional delay. When plotted against the hierarchical score difference (Fig.3C, red triangles), the area median functional delays exhibit strong correlation (r_p_=0.76 and P_p_<0.0001).

**Figure 3.**
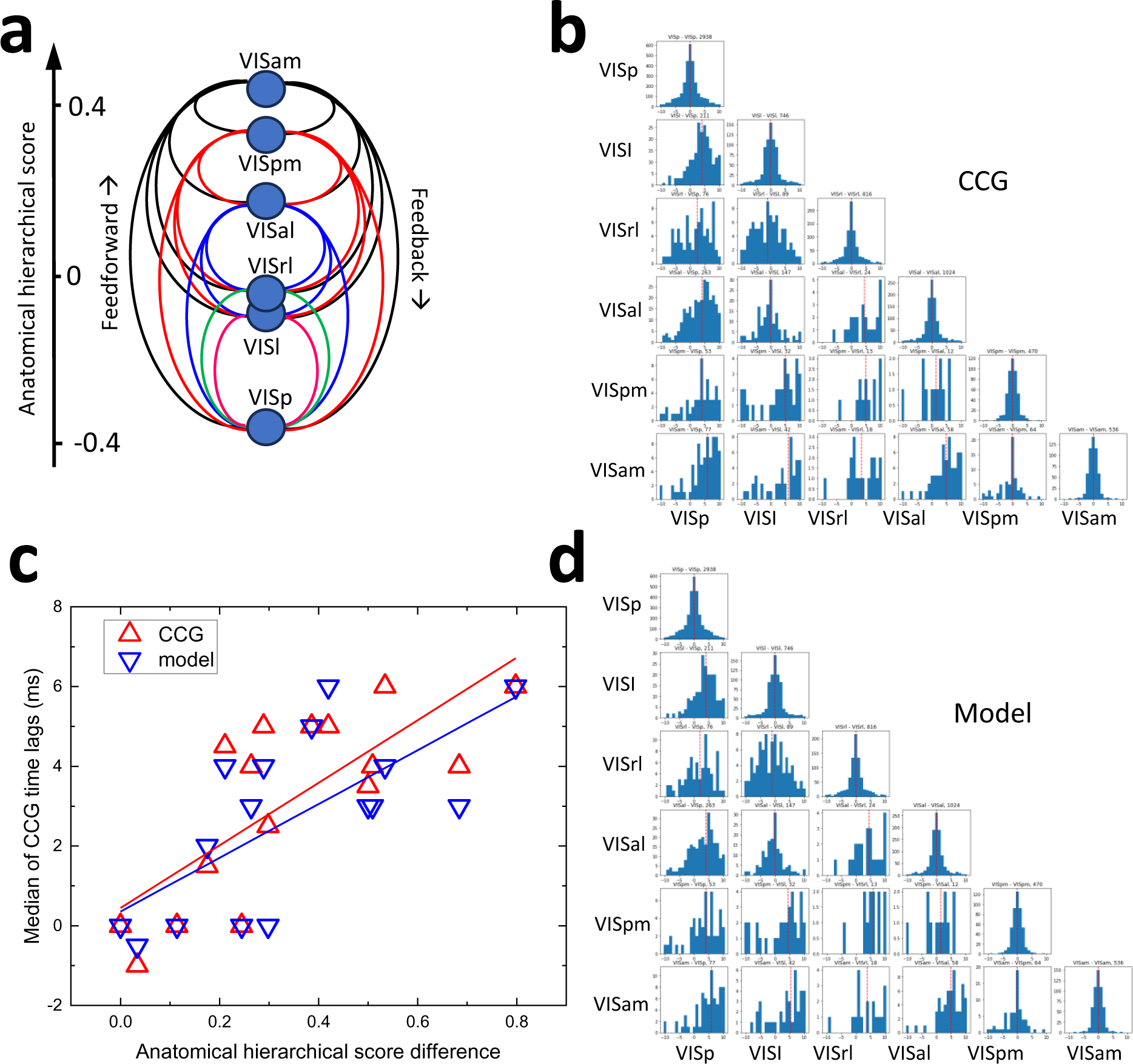
Static connectivity inferred by the model is biologically interpretable. **A)** Feedforward and feedback connections between 6 cortical areas ordered with respect to their anatomical hierarchy scores (after Ref.4). **B)** Histograms of the functional delays extracted from the jitter-corrected CCG for all pairs across all 26 sessions for each visual area combination. Dashed red line indicates the median of the distribution. **C)** Comparison of median functional delays vs anatomical hierarchy score difference extracted by DyNetCP (blue triangles) and by jitter-corrected CCG (red triangles) (VCN dataset, n=26 animals). **D)** Histograms of the functional delays inferred by static DyNetCP across all 26 sessions of VCN dataset for each visual area combination. Dashed red line indicates the median of the distribution.

The DyNetCP trained on the same 5023 pairs with **W**_**st**_ peak amplitudes above α_w_ of 7SD, whose shape is statistically indistinguishable (typical r_p_>0.9 and P_p_<0.0001) from the corresponding jitter-corrected CCG, produce similar functional delays (Fig.3D). The median functional delays predicted by DyNetCP by learning from all units (Fig.3C, blue triangles) also show strong correlation (r_p_=0.73, P_p_<0.0001) with the visual area anatomical hierarchical score difference^5^, very closely replicating (r_p_=0.99, P_p_<0.0001) the correlation observed with the descriptive pairwise CCG-based analysis. Therefore, the static DyNetCP is able to confirm previous observation of hierarchical organization of inter-area functional delays^15^.

### Comparison of inferred dynamic connectivity to descriptive statistical methods

Once the static connectivity **W**_**st**_ is established and verified, the dynamic stage of DyNetCP learns **W**_**dyn**_ that is strongly temporally modulated (Video 1). While no GT data are currently available to verify the model performance, model predictions can be compared to a classical pairwise Joint Peristimulus Time Histogram^39^ (JPSTH) that represents the joint correlations of spiking between two neurons and their variation across time. Specifically, given trial-aligned data with a trial length of *T* bins, the JPSTH is a *T* × *T* matrix where each entry (*a*, *b*) represents the percent of trials where neuron *i* spiked at time *a* and neuron *j* spiked at time *b*. For an example pair of excitatory spatially close units located in Layer 5 of the VISp area (session 766640955) corresponding JPSTH is shown in Fig.4A. Averaging over the columns of the JPSTH recovers the peristimulus time histogram (PSTH) of neuron *i* (left panel in Fig.3A), while averaging over the rows of the JPSTH recovers the PSTH for neuron *j* (bottom panel in Fig.3A). Since DyNetCP learns the conditional probability of the spiking of one neuron conditioned on others, we cannot directly compare dynamic weights to a standard JPSTH, which models joint correlations. Instead, for a proper comparison, we use a conditional joint peristimulus time histogram (cJPSTH), where we divide the columns of the JPSTH by the PSTH of neuron *i* (Fig.4B). This changes the meaning of each entry (*a*, *b*) in the cJPSTH that represents the conditional probability that neuron *j* spikes at time *b*, conditioned on neuron *i* having spiked at time *a* (Fig.4C).

**Figure 4.**
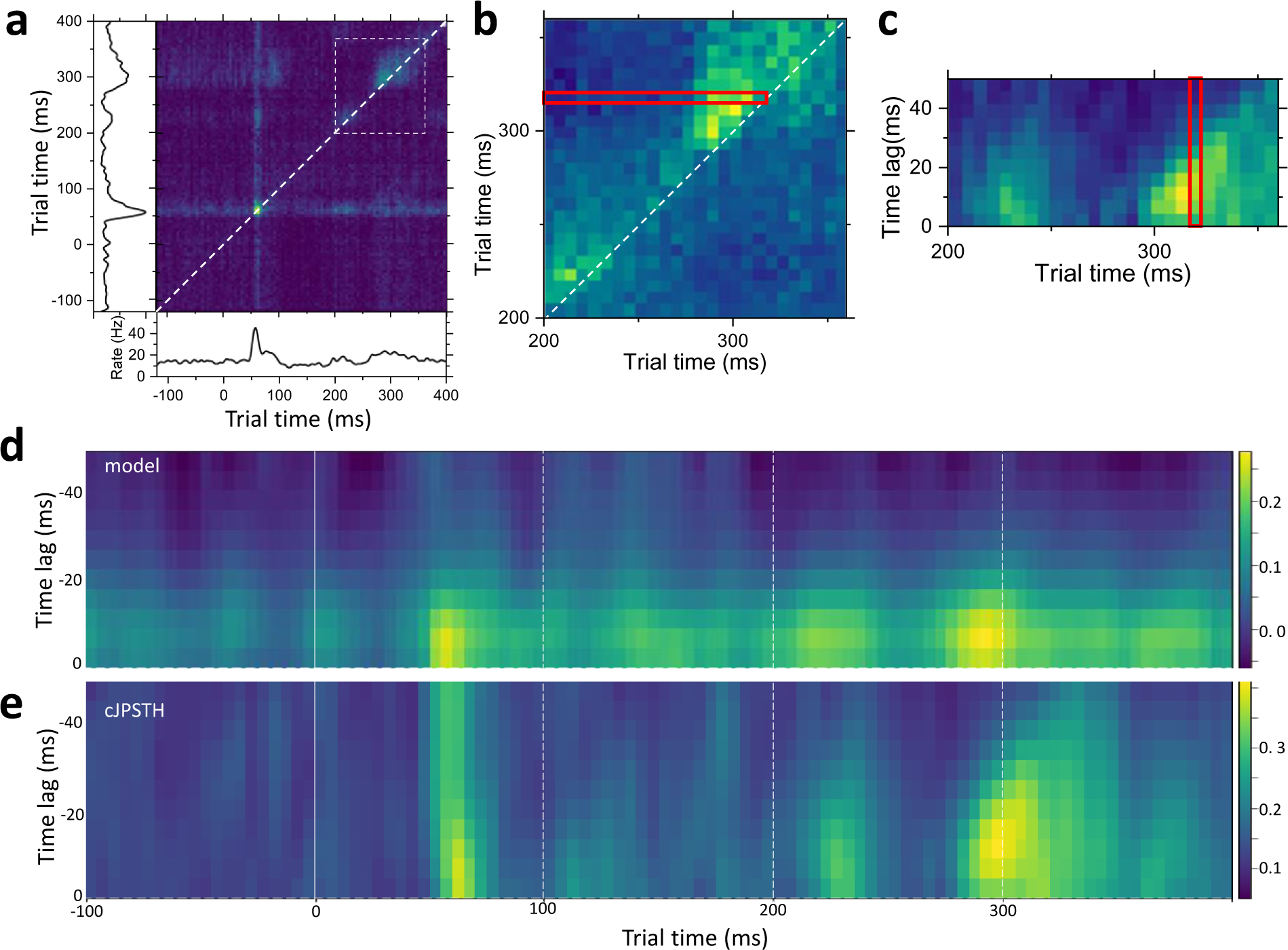
Comparison of inferred dynamic connectivity to descriptive statistical methods. **A)** An example of JPSTH for a local pair of excitatory units (session 766640955 of the VCN dataset, spatially close units from Layer 5 of the VISp area). Panels on the left and at the bottom show corresponding spike rates for receive and send units, correspondingly. White dotted square corresponds to the area of interest. See also Video 1. **B)** Construction of the conditional cJPSTH. Pixels from the white dotted square in A) are normalized by spiking rate of the send neuron. Row of pixels marked by red rectangle is used in C). **C)** cJPSTH corresponding to the white dotted square in A). Red rectangle corresponds to red rectangle pixels in B). **D)** Dynamic weight **W**_**dyn**_ for the same pair of excitatory units inferred for *L2* of 1 (see also Video 1). **E)** cJPSTH for the same pair of units.

**Fig.4 – figure supplement 1.**
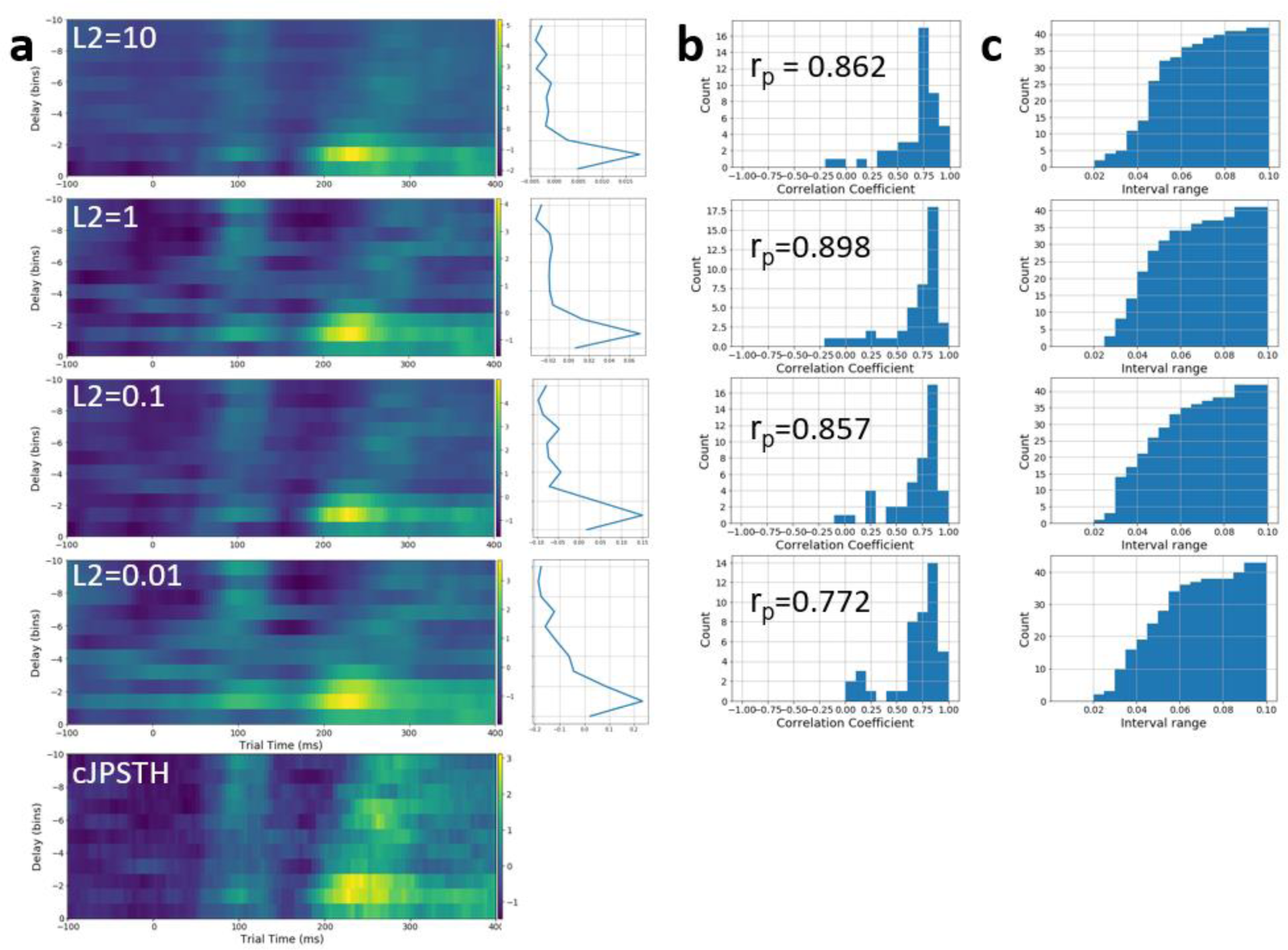
Tuning loss parameter. **A)** Set of dynamic weights **W**_**dyn**_ for a local pair of excitatory VISp neurons (session 771160300, same as in Fig.4D,E) learnt with different *L2* values. The corresponding cJPSTH is shown on the bottom. Left column shows a set of corresponding static weights **W**_**st**_. **B)** A histogram of Pearson correlation coefficients between the dynamic weights and cJPSTH for each pair (comparing each pixel of the weight/cJPSTH matrix) calculated for all 41 pairs found to be significant for this session. **C)** The two-sided paired equivalence tests (TOST) for all 41 pairs in this session indicate that the average difference between normalized weights inferred from both models is smaller than 0.09.

**Video 1:**
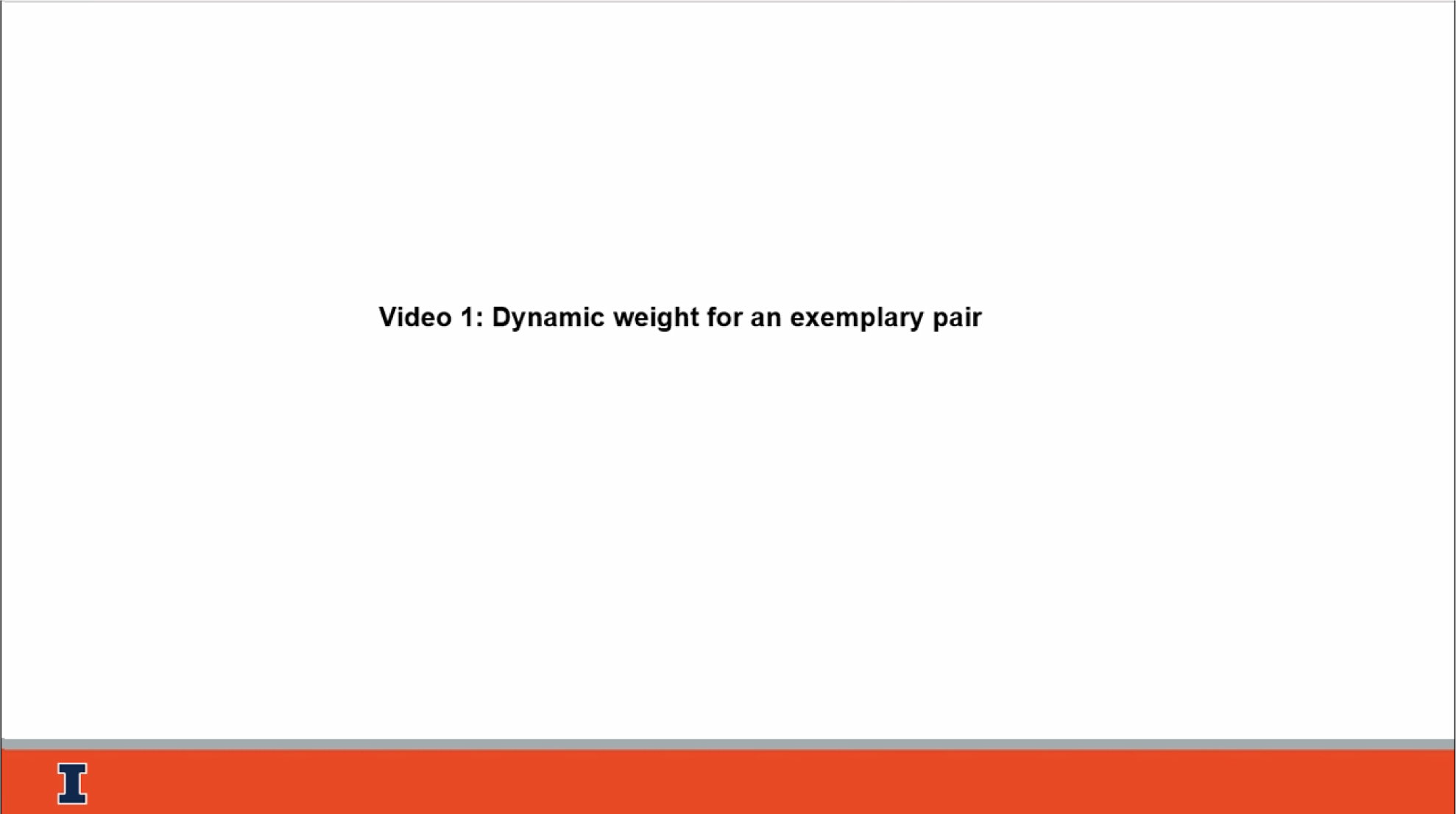
Dynamic weight for an exemplary pair. Dynamic weight **W**_**dyn**_ for a local pair of excitatory units from Layer 5 VISp area, session 766640955 (same pair as in Fig.4D, E). Static weight is shown on the background. Dashed horizontal line is a maximum (minimum) static weight. Static and dynamic weights are binned at 5ms. Available for download at [link](https://submit.biorxiv.org/submission/download?file=/hwpbp2/benchpress13/files/biorxiv/2023/552000/552512v4/supp/file00&msid=BIORXIV/2023/552512&roleName=author&gotoPageName=proof&continueToPage=proof&supp=true&request_journal_code=biorxiv&download_action_prefix=)

The dynamic weights **W**_**dyn**_ matrix inferred by DyNetCP for the same example pair (Fig.4D, see also (Video 1) reveal strong temporal modulation peaking at around 70ms, 220ms, 300ms, and 480ms indicating increased connection strength. Comparison with the cJPSTH matrix (Fig.4E) shows similar temporal variation. This comparison can be used to validate the choice of an optimal squared *L2* penalty that is introduced into the model loss function (Eq.6) to compensate the sparseness and irregularity of spiking often observed in cortical recordings (Methods). For a given set of thresholds α_*w*_, α_*s*_, and α_*p*_, scanning of *L2* from 0.01 to 10 (Fig.4 – figure supplement 1A) demonstrates statistically significant correlation (r_p_>0.77 and P_p_<0.001) of the dynamic weights **W**_**dyn**_ matrices and cJPSTH matrices observed for all *L2* values with the best results obtained for *L2* between 0.1 and 1 (Fig.4 – figure supplement 1B). The two-sided paired equivalence test (TOST) (Fig.3 – figure supplement 1C) confirms that the average difference between normalized weights inferred from both models is smaller than 0.09. Therefore, the DyNetCP reliably recovers the time-dependent dynamic weights **W**_**dyn**_ even in the presence of strong nonstationary spiking irregularities and sparse spiking.

### Model enables discovery of cellular-level dynamic connectivity patterns in visual cortex

The model architecture (Fig.1A) designed with two interacting static and dynamic parts enables justification of the choice of model parameters and verification of model inference results by comparison with the results of descriptive statistical methods. Therefore, even in the absence of experimental or synthetic datasets with known ground truth (GT) of cellular-level dynamic connectivity, the model can provide an otherwise unattainable insights into dynamic flows of information in cortical networks. To ensure statistical significance of inferred connections three thresholds (Methods) were applied simultaneously to each of the 26 sessions in the VCN database: a unit average spike rate α_*s*_>1Hz, a pair spike rate threshold α_*p*_>4800, and a static weight peak threshold α_*w*_>7SD that produced 808 valid units forming 1020 valid pairs across all 6 cortical areas (Methods, Table 1).

### Reconstruction of dynamic connectivity in a local circuit

To illustrate reconstruction of static and dynamic connectivity in a local circuit we use recording from a single animal (session 819701982) from a single Neuropixel probe implanted in the VISam area. Application of three thresholds α_*s*_, α_*p*_, and α_*w*_ to all clean units passing quality metrics (Methods) produce 6 pairs with statistically significant (α_*w*_>7SD) and narrow (<2ms) peak in a static weight **W**_**st**_ (Methods, Table 2) indicating strong putative connections (Fig.5A). Units are classified as excitatory (waveform duration longer than 0.4ms) or inhibitory (less than 0.4ms) and are assigned to cortical depth and cortical layers based on their spatial coordinates (Methods). Directionality of the connection is defined by negative functional time lag in a static weight **W**_**st**_ (or jitter corrected CCG) binned at 1ms. Local circuit reconstructed from static weights matrices is shown in Fig.5B. Corresponding dynamic matrices **W**_**dyn**_ (Fig.5C) for the same pairs demonstrate strong temporal modulation (see also Video 2). Prior to presentation of a stimulus (t<0ms) the inhibitory drives (negative values of **W**_**dyn**_ marked by deep blue color in Fig.5C) prevail. The first positive peak of **W**_**dyn**_ around 70ms (positive values of **W**_**dyn**_ marked by bright yellow color in Fig.5C) is shared by all but one pair. Such synchronous spiking at short time lags does not necessarily indicate a TP direct connection but can also be an FP artefact induced by arrival of afferent signals^44^. However, at later times, additional strong peaks appear for some of the pairs that are not accompanied by corresponding increase of the firing rates. Some of the pairs form an indirect serial chain connections (e.g., 3→2 and 2→4 Fig.5B) that can also generate confounding FP correlations that is notoriously difficult to identify using descriptive statistical models^24^. However, the DyNetCP being a probabilistic model might lessen the impact of both the common input and the serial chain FP artefacts by abstracting out such co-variates that do not contribute significantly to a prediction. For example, the **W**_**dyn**_ for a potential 3→4 artefactual connection does not show peaks expected from the serial chain.

**Figure 5.**
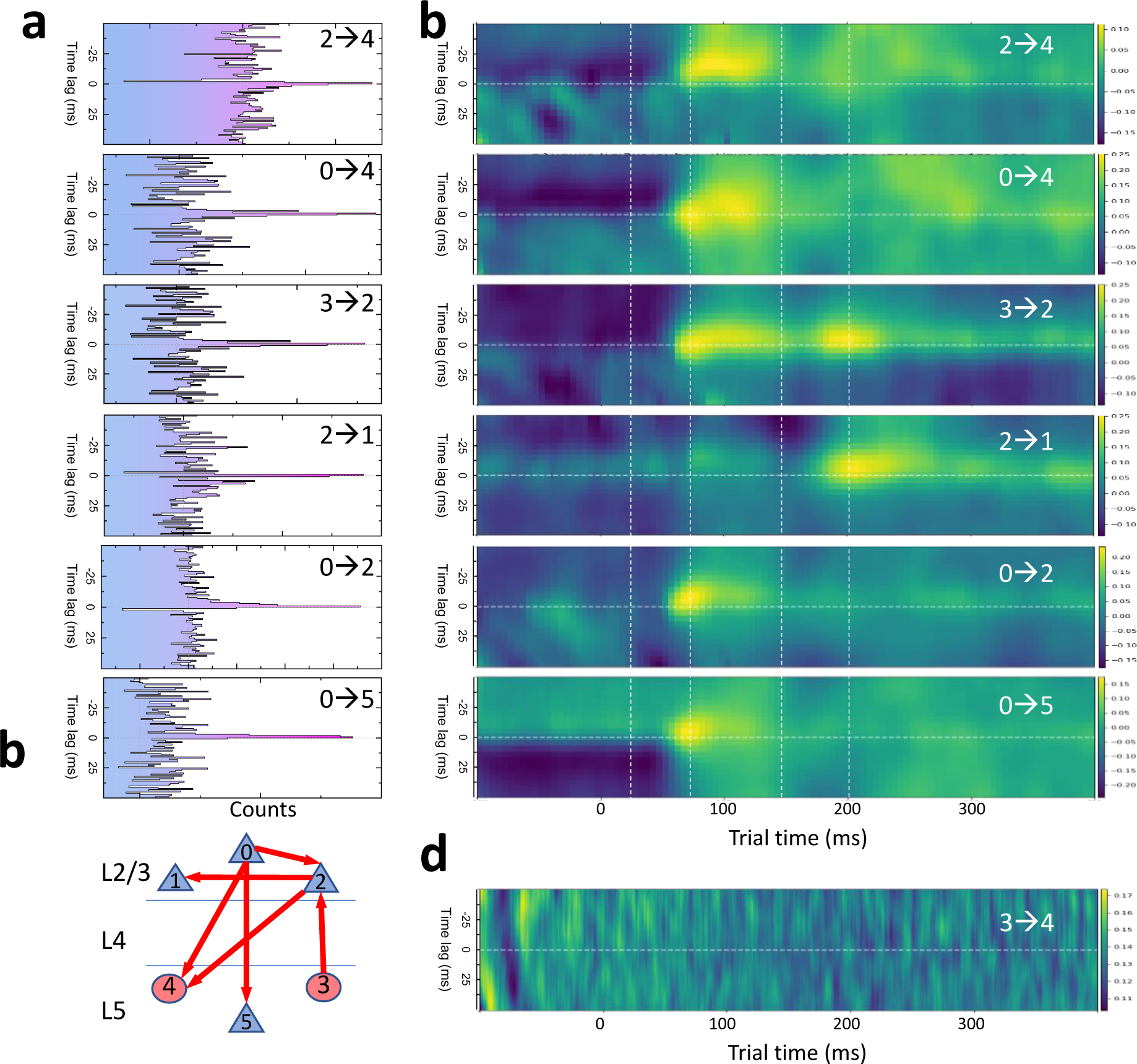
Model enables discovery of cellular-level dynamic connectivity patterns in visual cortex. **A)** Jitter-corrected static weights **W**_**st**_ for a connected pairs in an example local circuit (session # 819701982, VISam area). Units are numbered by local indices (Methods, Table 2). **B)** Dynamic weights **W**_**dyn**_ for the same pairs. **C)** Schematic diagram of static directional connectivity in pairs in a local circuit in A). Circles and triangles correspond to excitatory and inhibitory units, respectively. **D)** An example of a dynamic weight **W**_**dyn**_ for a pair (3→4) classified as un-connected.

**Video 2:**
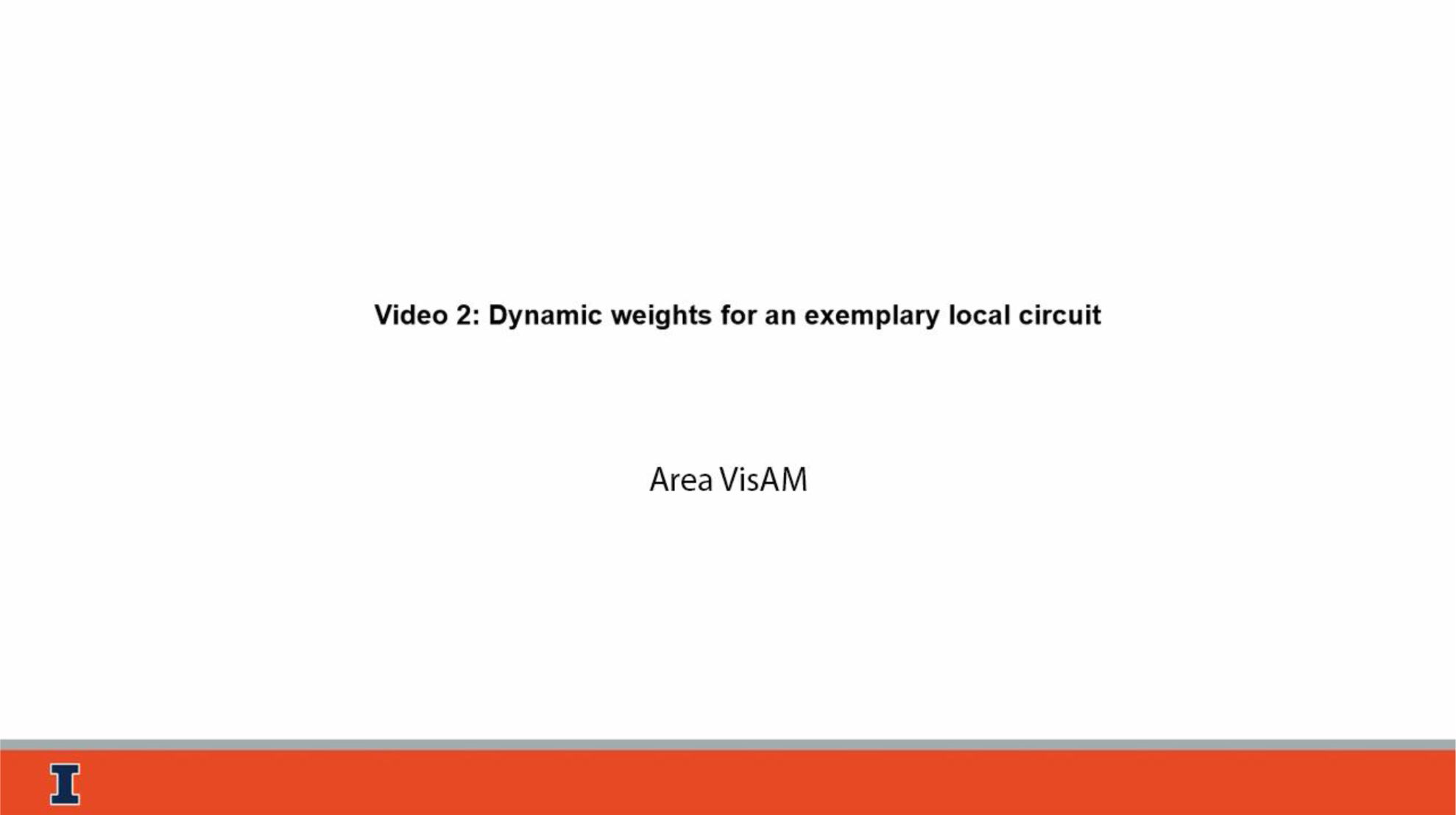
Dynamic weights for an exemplary local circuit. Top row: Dynamic connectivity for an example local circuit (same as in Fig.5) from VISam area, session 819701982 for different trial times. Circles and triangles correspond to excitatory and inhibitory units, respectively. Units are numbered by local indices and ordered with respect to cortical layers. The size of blue (red) arrows reflects the dynamic weight of the corresponding connection at a given time. Bottom two rows: Dynamic weight **W**_**dyn**_ for each pair in the circuit. Dashed horizontal line is a maximum static weight. Static and dynamic weights are binned at 5ms. Available for download at [link](https://submit.biorxiv.org/submission/download?file=/hwpbp2/benchpress13/files/biorxiv/2023/552000/552512v4/supp/file01&msid=BIORXIV/2023/552512&roleName=author&gotoPageName=proof&continueToPage=proof&supp=true&request_journal_code=biorxiv&download_action_prefix=)

**Table. 2.**
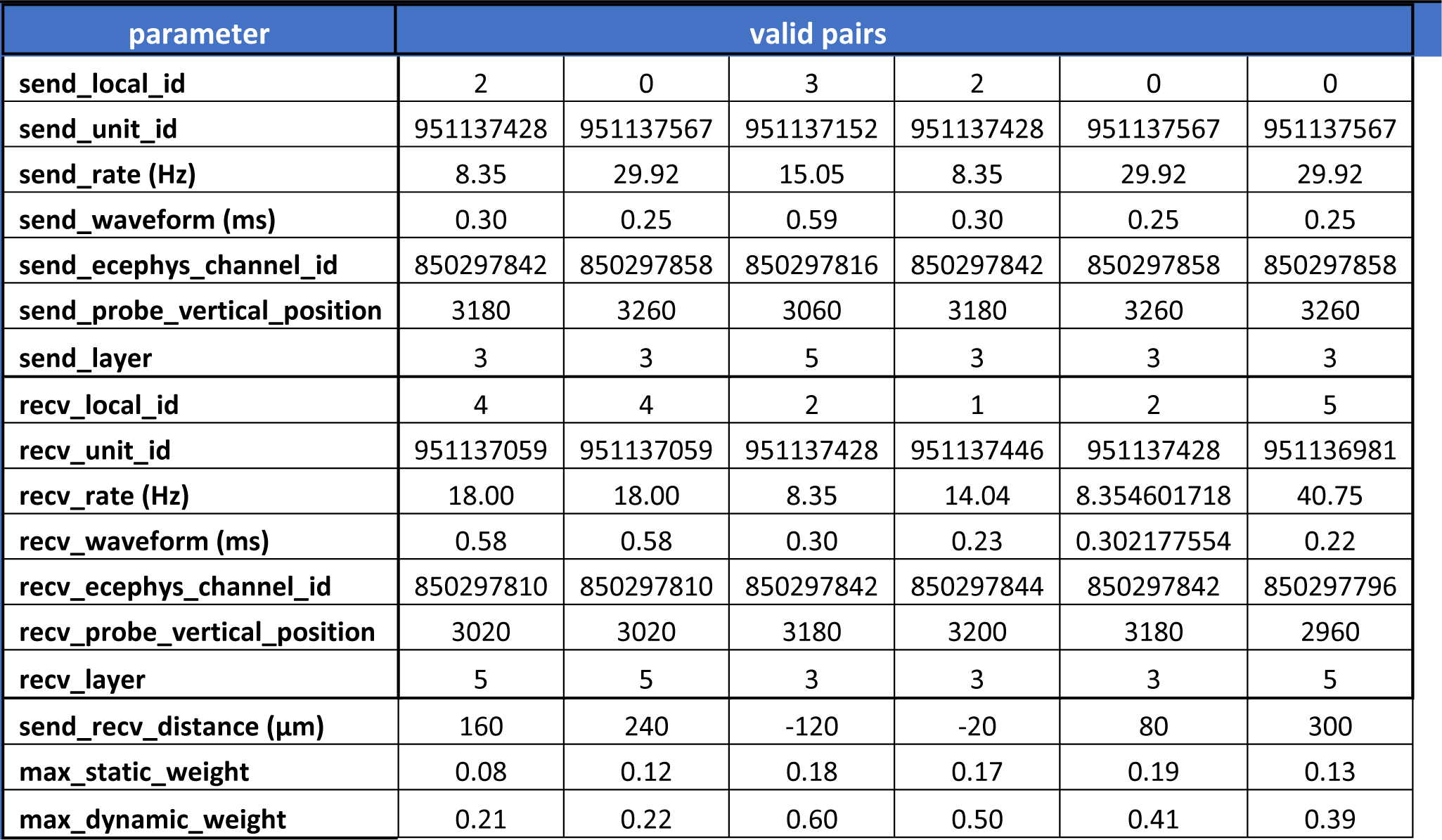
Experimental dataset used for reconstruction of local circuit in Fig.5 and Fig.6. Clean units (passed unit quality metrics) that from valid pairs (passed through all three thresholds α_*s*_, α_*p*_, and α_*w*_) from recordings in VISam area in session 819701982. Units are classified as excitatory (waveform duration longer than 0.4ms) or inhibitory (less than 0.4ms) and are assigned to cortical depth and cortical layers based on their spatial coordinates. Local circuit reconstructed from static weights matrices (max_static weight) is shown in Fig.5B. Maximum dynamic weight (max_dynamic_weight) is taken at functional time lag. Distance between units (send_recv_distance) is estimated from vertical positions of corresponding principal electrodes (probe_vertical_position).

To aid identification of such confounding FP connections, we performed preliminary tests on a synthetic dataset that contain a number of common input and serial chain triplets and were able to identify them by comparing **W**_**dyn**_ values at a given trial time to corresponding **W**_**st**_ (Methods). Figure 6 shows temporal evolution of normalized dynamic weight values taken from Fig.5C along the trial time axis at corresponding functional time lags. We assign excitatory (inhibitory) connection to a given pair as present when the ratio **W**_**dyn**_⁄**W**_**st**_ is larger (or smaller) than 1 (−1). Corresponding dynamic graphs for a local circuit of Fig.5 are shown in Fig.6B for characteristic trial times. Such dynamic circuit reconstruction is aided by observation of the troughs (inhibitory connection) and peaks (excitatory connection) in the **W**_**dyn**_ cross-sections along the lag time axis taken at a given trial time (Fig.6 – figure supplement 1, see also Video 2).

**Fig. 6.**
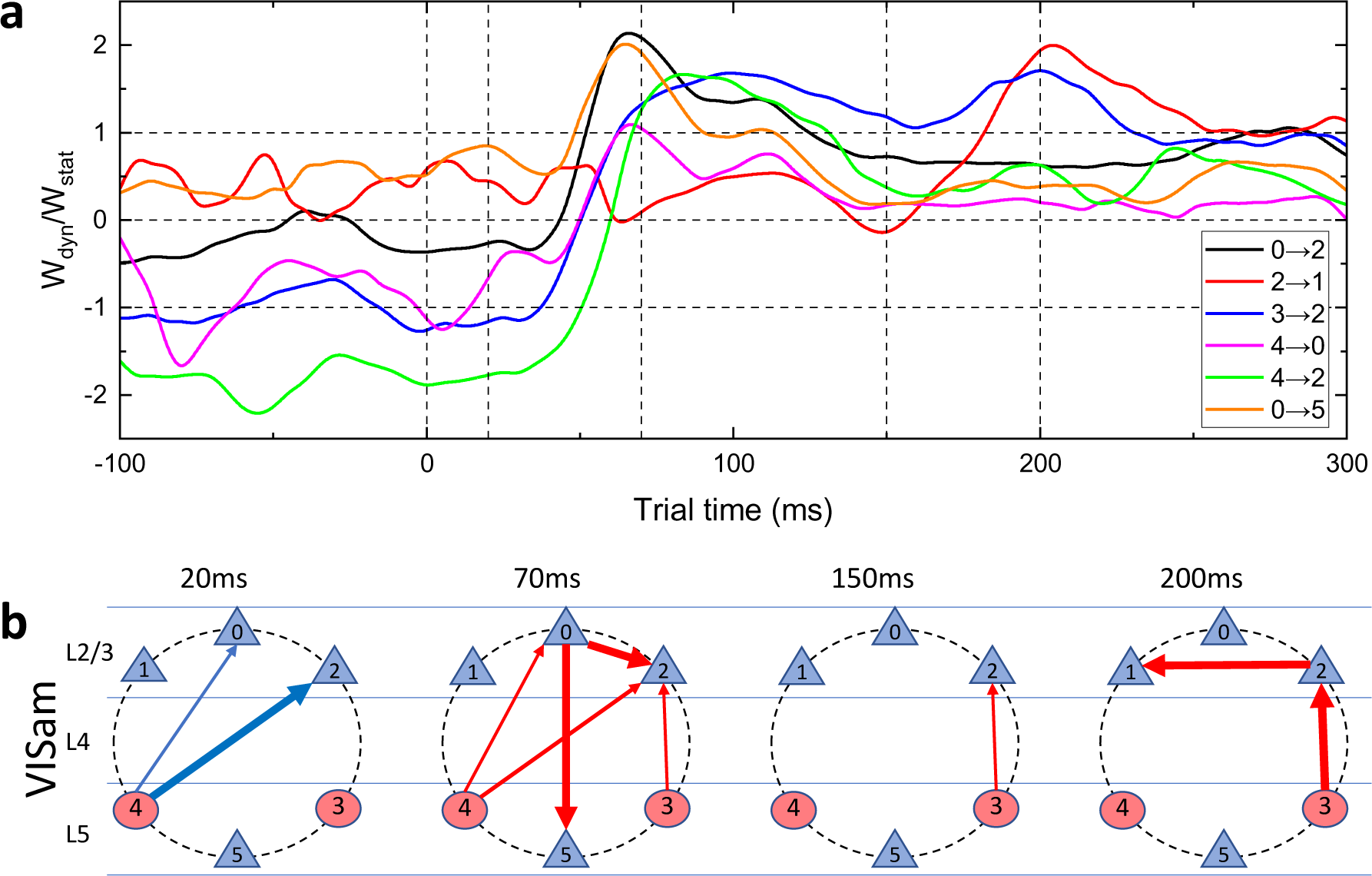
Temporal variation of dynamic weight and reconstruction of dynamic local circuit. **A)** Normalized dynamic weights for all valid pairs in an example local circuit (session # 819701982, VISam area). Units are numbered by local indices. **B)** Schematic diagram of dynamic connectivity in a local circuit for different trial times. Circles and triangles correspond to excitatory and inhibitory units, respectively. The size of blue (red) arrows reflects the dynamic weight of the inhibitory (excitatory) connection at a given time.

**Figure 6 - figure supplement 1.**
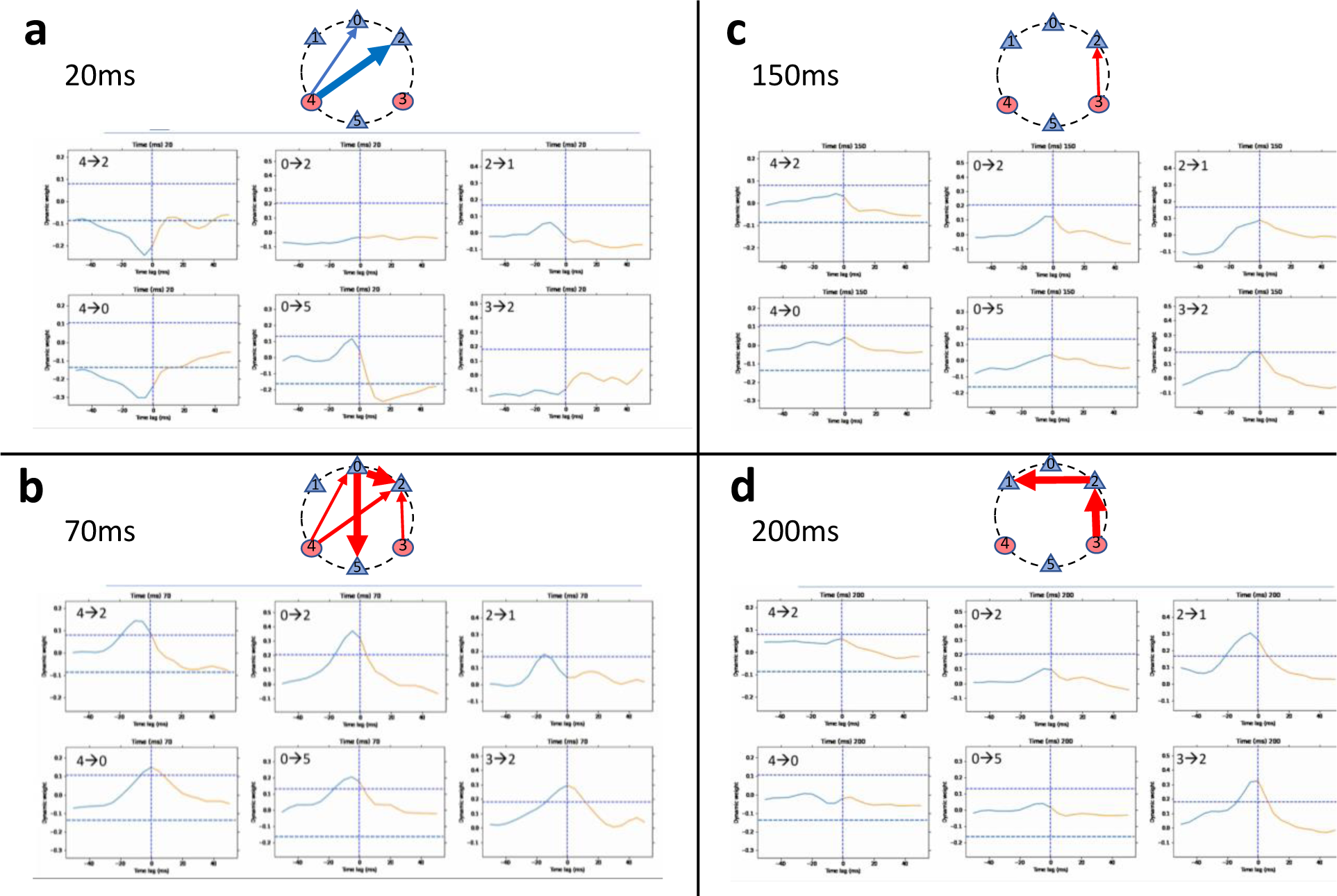
Reconstruction of a dynamic connectivity in a local circuit. Dynamic weights **W**_**dyn**_ for 6 valid pairs in an example local circuit in Fig.6 taken at different trial times. **A)** at 20ms. **B)** at 70ms. **C)** at 150ms. **D)** at 200ms. Reconstruction of dynamic connectivity is shown on top of each figure. Circles and triangles correspond to excitatory and inhibitory units, respectively. The size of blue (red) arrows reflects the dynamic weight of the inhibitory (excitatory) connection at a given time. See also Video 2.

**Video 3:**
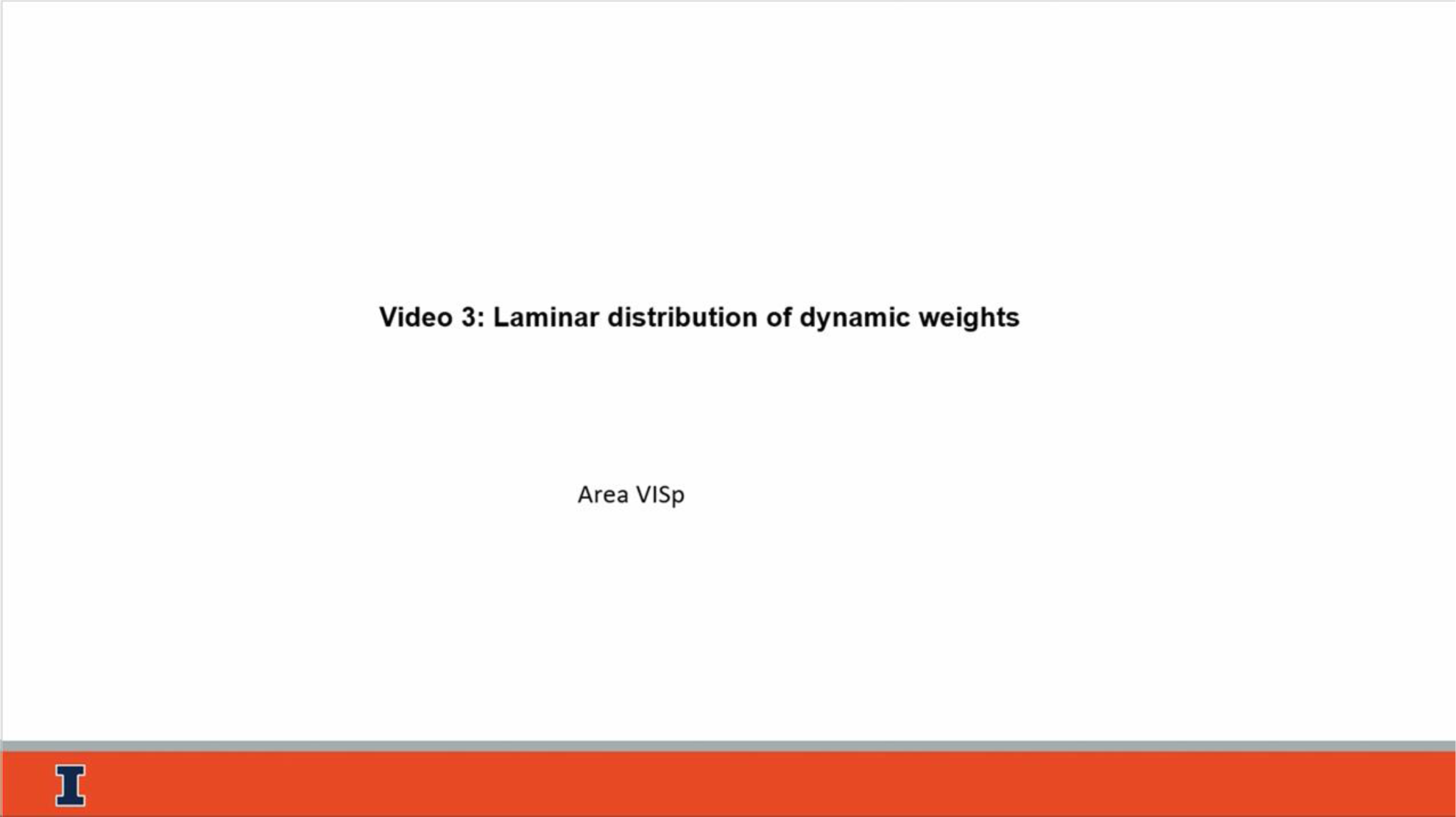
Laminar distribution of dynamic weights for VISp area. **W**_**off**_ for a VISp area for all n=26 animals (same as in Fig.7A) for different trial times. Units ordered by cortical depth and assigned to cortical layers. The circles diameter is proportional to the dynamic offset. Red, blue, and magenta circles correspond to excitatory-excitatory, inhibitory-inhibitory, and mixed connections, respectively. Available for download at [link](https://submit.biorxiv.org/submission/download?file=/hwpbp2/benchpress13/files/biorxiv/2023/552000/552512v4/supp/file02&msid=BIORXIV/2023/552512&roleName=author&gotoPageName=proof&continueToPage=proof&supp=true&request_journal_code=biorxiv&download_action_prefix=)

Note, that since DyNetCP, as opposed to population dynamic models^26–29^, learns latent variables assigned to individual units, the anatomical and cell-type specific information in static graph of Fig.5B is preserved. As in Fig.5B, units in the circuit of Fig.6B are ordered with respect to their cortical depth, assigned to specific cortical layers, classified as inhibitory (excitatory) based on their waveform duration. Further analysis of optotagging^45^ (Methods) was used to identify that none of inhibitory units in this local circuit belong to a Vasoactive Intestinal Polypeptide (Vip) cell class. DyNetCP can also be trained separately on trials when animal was running, and animal was at rest (Methods, Table 1) to explore the role of brain state on dynamic connectivity patterns. Therefore, not only DyNetCP can provide static connections in this local circuit, but, more importantly, can also track the time epochs when individual cellular-level connections are activated or inhibited that are time-aligned to the stimulus presentation, brain state, and behavior.

### Dynamic connectivity in VISp primary visual area

Going beyond local circuits, the dynamic analysis of information flows during stimulus presentation can be extended to area-wide dynamic maps. However, with the aggressive choice of threshold parameters (α_*s*_>1Hz, α_*p*_>4800, α_*w*_>7SD) the average number of valid pairs per session (Methods, Table 1) is relatively low to generalize (mean:39 ± 4 SEM). To analyze area-wide connectivity, an example of cellular-level dynamic interactions in primary visual area VISp learned from all 26 sessions is shown in Video 3. Still frames from this video are presented in Fig.7A for a number of characteristic trial times. Units are ordered by cortical depth and assigned to cortical layers. Circles represent the magnitude of a dynamic weight **W**_**dyn**_ taken at corresponding functional time lag. The circles diameter is proportional to the dynamic offset **W**_**dyn**_ − **W**_**st**_. The dynamic weights are changing over the time of the trial (Video 3) and exhibit interactions between excitatory cells (red circles), inhibitory cells (blue circles) and mixed drives (magenta circles). While excitatory-excitatory connections are mostly activated within layer 4 and layer 5, connections involving inhibitory cells dominate inter-layer interactions between all the layers. Feedforward extra-laminar interactions (e.g., input layer 4 is driving supergranular layer 2/3 and infragranular deep layers 5 and 6) reveal formation of neuronal ensembles across all the cortical depth. Feedback interactions are also distributed across the cortical depth with nearly the same maximum weights. These observations are inconsistent with static connection efficacy extracted from CCG peaks recorded in V1 columns in anesthetized monkeys^46^ that indicate the efficacy greatly diminished within <200μm from the source. It is consistent, however, with static spike transmission strength extracted from CCG peaks in V1 columns in mice^47^ that reveal much strong increase of long-distance correlations between layer 2/3 and layer 5 during waking in comparison to sleep states. In general, not only brain state can modulate the connection strength, but most importantly, as shown in Video 3 and Fig.7A, the connection strength between all the layers is strongly temporally modulated during the stimulus presentation. Therefore, our results demonstrate that tracking dynamic connectivity patterns in local cortical networks can be invaluable in assessing circuit-level dynamic network organization.

**Figure 7.**
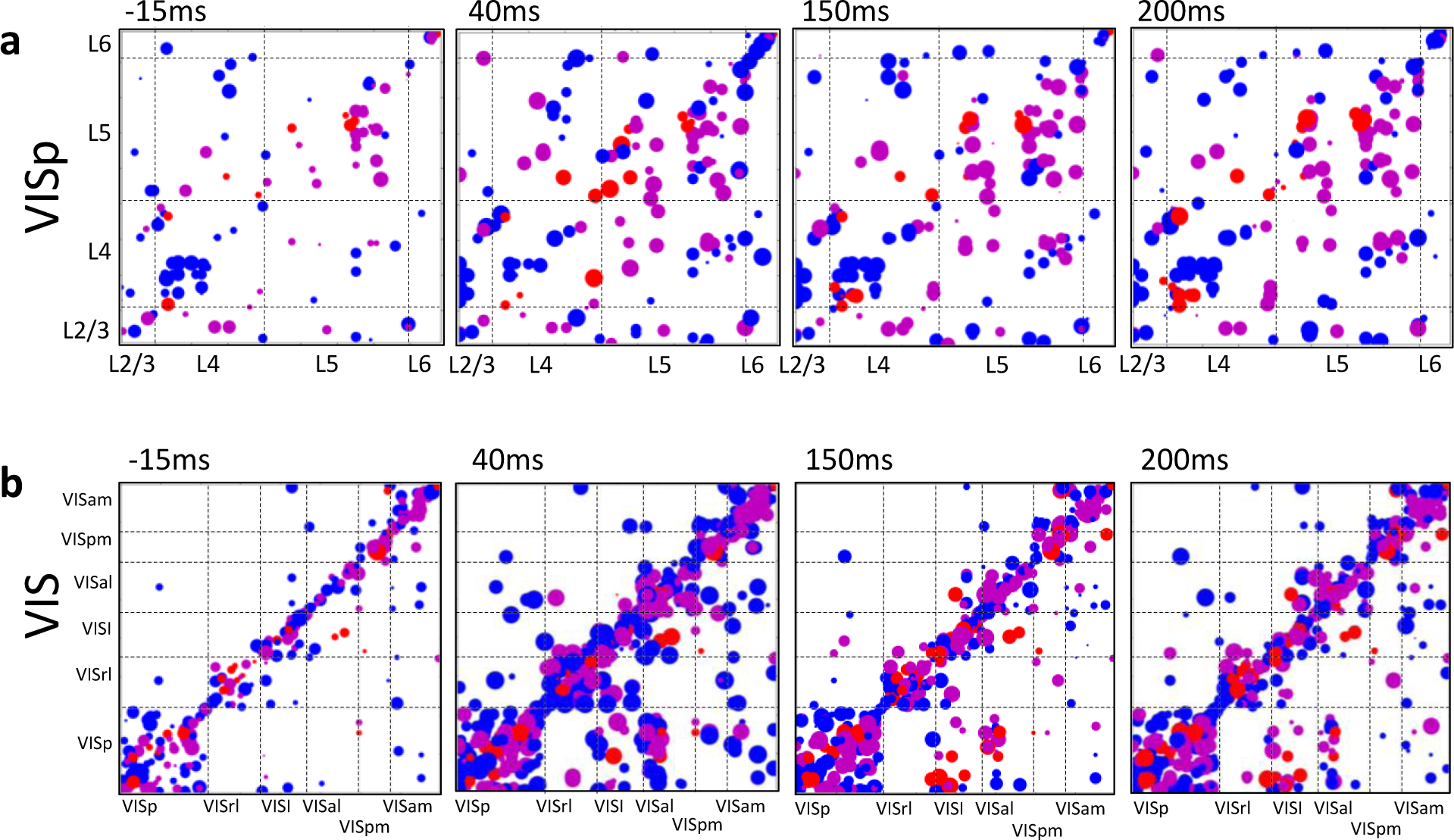
Dynamic functional connectivity prediction along the hierarchy axis of visual cortices. **A W**_**off**_ at different trial times (VISp area, n=26 animals). Units ordered by cortical depth and assigned to layers. The circles diameter is proportional to the dynamic offset. Red, blue, and magenta circles correspond to excitatory-excitatory, inhibitory-inhibitory, and mixed connections, respectively. **B) W**_**off**_ matrices for all visual areas (n=26 animals) ordered by their hierarchical

**Video 4:**
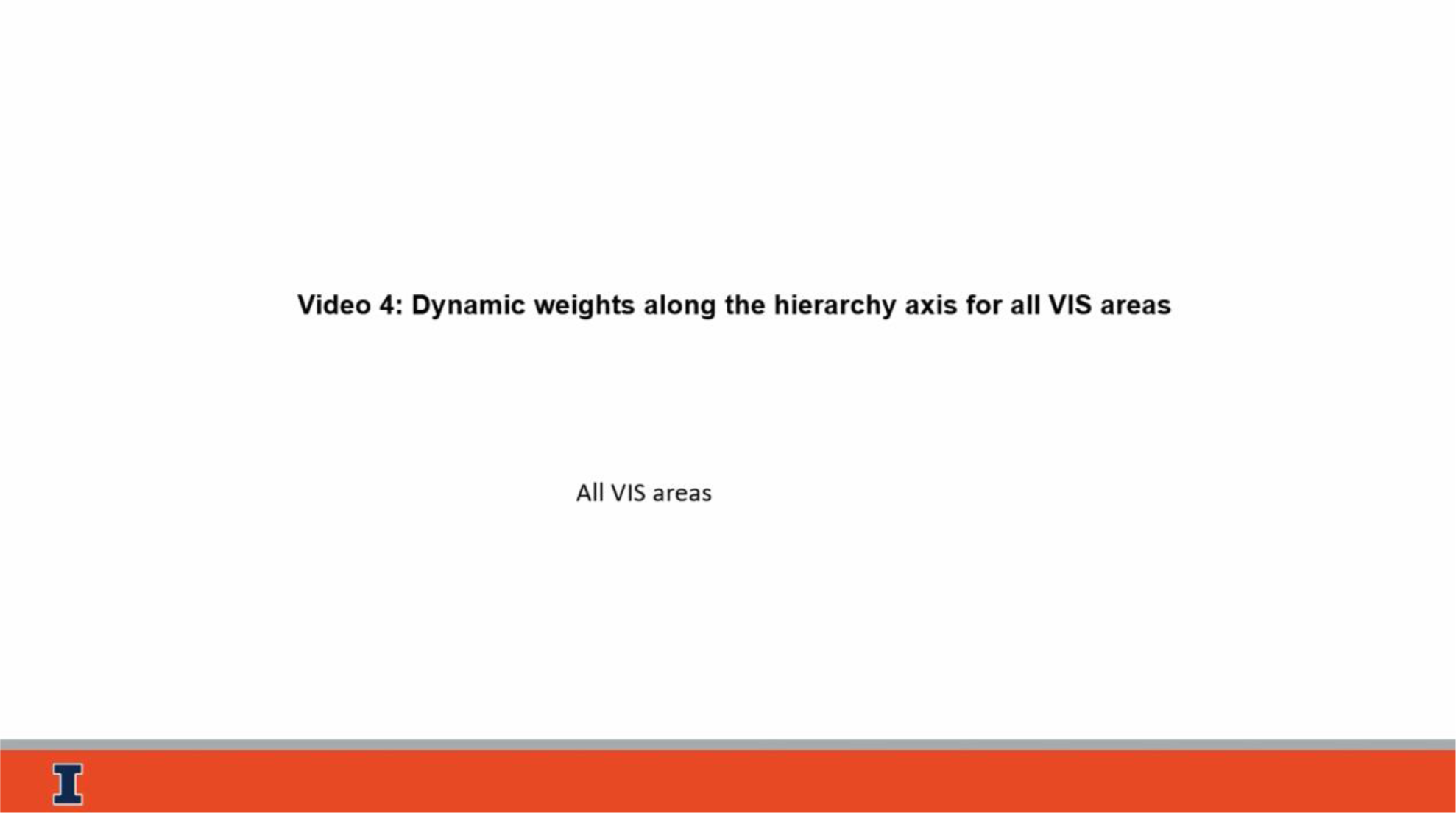
Dynamic weights along the hierarchy axis for all VIS areas. **W**_**off**_ for all VIS areas for all n=26 animals (same as in Fig.7B). Units ordered by cortical depth and assigned to cortical layers. The circles diameter is proportional to the dynamic offset. Red, blue, and magenta circles correspond to excitatory-excitatory, inhibitory-inhibitory, and mixed connections, respectively. Available for download at [link](https://submit.biorxiv.org/submission/download?file=/hwpbp2/benchpress13/files/biorxiv/2023/552000/552512v4/supp/file02&msid=BIORXIV/2023/552512&roleName=author&gotoPageName=proof&continueToPage=proof&supp=true&request_journal_code=biorxiv&download_action_prefix=)

### Dynamic connectivity along the hierarchy axis of all visual cortices

Similarly, cellular-level dynamic interactions at the cortex-wide level for all 6 cortical areas learned from all 26 sessions are shown in Video 4. Still frames from this Video for the same characteristic times are shown in Fig.7B. Inter-areal long-range interactions across various visual cortices (off-diagonal terms in Fig.7B) exhibit complex temporal evolution with rich feedforward (along the anatomical hierarchical score difference^5^) and feedback interactions. Therefore, DyNetCP can uncover the dynamics of feedforward- and feedback-dominated epochs during sensory stimulation, as well as behavioral states (e.g., epochs in the VCN dataset when animals are running or sitting), however, as opposed to population decoding models^26–29^, produce cellular-level cell-type specific connectivity patterns that are otherwise hidden from analysis.

## DISCUSSION

DyNetCP can reveal cellular-level dynamic interactions within local circuits, within a single brain area, as well as across global hierarchically organized information pathways. Such detailed information, although present in the original datasets, is otherwise hidden from usual analysis that typically relies on time-averaged spike rate coding or population-averaged coding.

Biological interpretation of the extracted dynamic weights can follow the terminology of the short-term plasticity between synaptically connected neurons^25,33–37^ or spike transmission strength^30–32,47^. Alternatively, temporal changes in connection weights can be interpreted in terms of dynamically reconfigurable functional interactions of cortical networks^8–11,13,46^ through which the information is flowing. We could also not exclude interpretation that combines both ideas. In any event our goal here is to extract these signals for a pair (video1, Fig.4), a cortical local circuit (Video 2, Fig.5), and for the whole visual cortical network (Videos 3, 4 and Fig.7). We envision that the next step here, falling beyond the scope of the current paper, would be to correlate the time dependence of the extracted dynamic weights with the stimulus presentation and the dynamics of the behavioral parameters during the decision-making process to infer where and how the information is flowing in the timescale of a decision making.

It is instructional to assess whether the temporal cellular-level interactions uncovered by DyNetCP are bringing new latent dimensions to the analysis of neural code. Dimensionality reduction methods^18,43,48^ are often used to estimate the number of independent dimensions of the code that can better explain the variability of spike trains. When principal component (PC) dimensionality reduction^29^ is applied to the dynamic weights **W**_**dyn**_ of all valid pairs (i.e., passed through all 3 thresholds) in a single animal (65 units, 105 pairs, session 831882777), up to 33 components are required to account for 95% of the variance (Fig.8A, black). In contrast, only 5 PC are sufficient to explain 95% variance of spike rates for the same units (Fig.8A, red). If all clean units in all visual cortices are taken (264 VIS units) the dimensionality increases, but only slightly to 8 PC (Fig.8A, blue). The normalized intrinsic dimensionality (NID) estimated as a number of major PC components divided by the number of units is between 0.03 and 0.08 for spike rates exhibiting striking simplicity consistent with an often observed trend^49^. However, in contrast, the NID for dynamic weights is as large as 0.5 showing dramatic increase of dimensionality.

**Figure 8.**
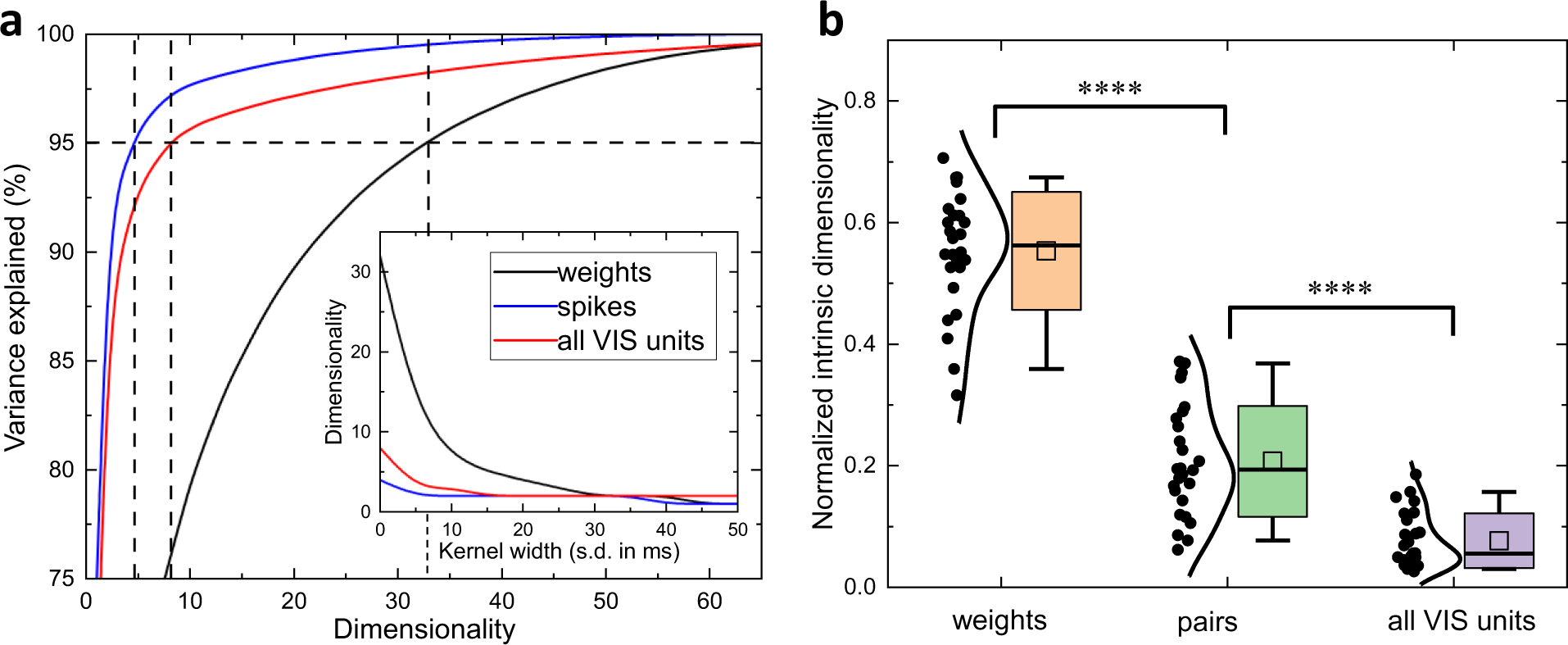
Intrinsic dimensionality of dynamic connectivity is large. **A)** Intrinsic dimensionality of neural activity manifold (session 831882777, no temporal smoothing) using spiking data for all VIS cortical units (red), units in selected pairs (blue), and for the same unit pairs using dynamic weights (black). **Inset**: Intrinsic dimensionality as a function of temporal gaussian kernel width. B) Intrinsic dimensionality (n=26 animals) in spike rate space normalized by the number of all VIS units (violet) and units in selected pairs (green). Intrinsic dimensionality in a weight space for units in selected pairs (orange). Box: SD, whiskers: 5%-95% range, box: mean, horizontal line: median, distribution: kernel Scott, **** is p-value<0.0001.

This result holds for each of the 26 sessions with the NID in the spiking rate space for valid pairs (Fig.8B, green, mean:0.20±0.018 SEM) and for all VIS units (Fig.8B, violet, mean:0.08±0.01 SEM). In contrast, much larger NID is observed in the space of dynamic weights (Fig.8B, orange, mean:0.55±0.02 SEM) indicating that the dimensionality of a dynamic network manifold is becoming comparable to the number of recorded units. Hence the observed complexity stems from a relatively small subpopulation (valid units in pairs mean: 42.7±3.3 SEM) comprising just 20% of the total (all clean VIS units mean: 194.9±11.9 SEM) and at relatively short time scales below 30ms when it survives temporal Gaussian kernel smoothing (inset in Fig.8A). DyNetCP, therefore, reveals the dynamically reconfigurable functional architecture of cortical networks that is otherwise hidden from analysis.

In summary, DyNetCP provides a fully reproducible and generalizable processing framework that allows investigators to explore often overlooked dimensions of the neural code that are potentially rich for novel biological insights. We believe that our approach can help to provide a conceptual link between brain-wide network dynamics studied with neuroimaging methods and modern electrophysiology approaches resolving such dynamics at a cellular level.

## METHODS

### 1. Model implementation and training

#### Computational resources

The DyNetCP model can be trained in a single step to produce both static and dynamic weight matrices simultaneously. Alternatively, static, and dynamic training can be done sequentially to minimize the computational resources and to enable intermediate checking of the model parameters with respect to descriptive statistical results. In this case, prior to implementing the dynamic part of DyNetCP, the pairs are classified as putative connections based on a statistical analysis (see below) of the static weights, and the dynamic component is trained only on these pairs.

Computation time and memory consumption can also be optimized by the choice of the bin size *d* at the cost of temporal precision of the weight prediction. To better identify putative connections and to ease comparison with descriptive CCG models, we use a bin size of 1ms for training of static weights. For training the dynamic part, we use spiking data binned at a 5ms resolution. Evaluation of statistical significance of the model predictions (see below) indicates that a 5ms bin size strikes a good balance between computational efficiency, temporal resolution, and data sparsity.

All models are trained using the ADAM optimizer^50^, a batch size of 16, and a learning rate of 0.0001 for 2000 epochs, where the learning rate is reduced by a factor of 10 after 1000 epochs. For each dataset, we set aside some of the data for validation. We use the validation data for early stopping: after every epoch of training, we compute the loss on the validation data, and terminate training early if the validation loss has not improved for a specific number of epochs (100 when training the static portion and 50 when training the dynamic portion). Note, to properly assess generalization, all plots in this work show DyNetCP static and dynamic weights outputs from the validation data.

When training the static part of the model on the synthetic HH dataset^25^, we use λ = 1. For training the static part on the VCN dataset^15^ we use λ = 100, which we found to lead to a lower residual when comparing to CCG values. For training the dynamic weights for all models we use λ = 10.

#### Averaging Dynamic Weights

To extract a strong signal from the DyNetCP dynamic weights, we need to aggregate information over experimental trials. However, we do not want to simply average the weights over trials, as all weights which are not paired with an incoming spike (i.e., all weights 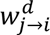 such that 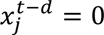) do not contribute to the spike rate prediction and therefore are not constrained to capture the spiking correlations. As a result, when computing trial-aggregated DyNetCP weights, we average over weights that are time-aligned with spike times of corresponding unit (Fig.9A) as follows:

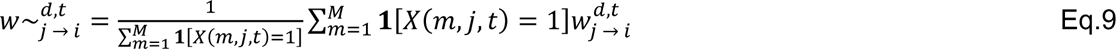

**Fig. 9.**
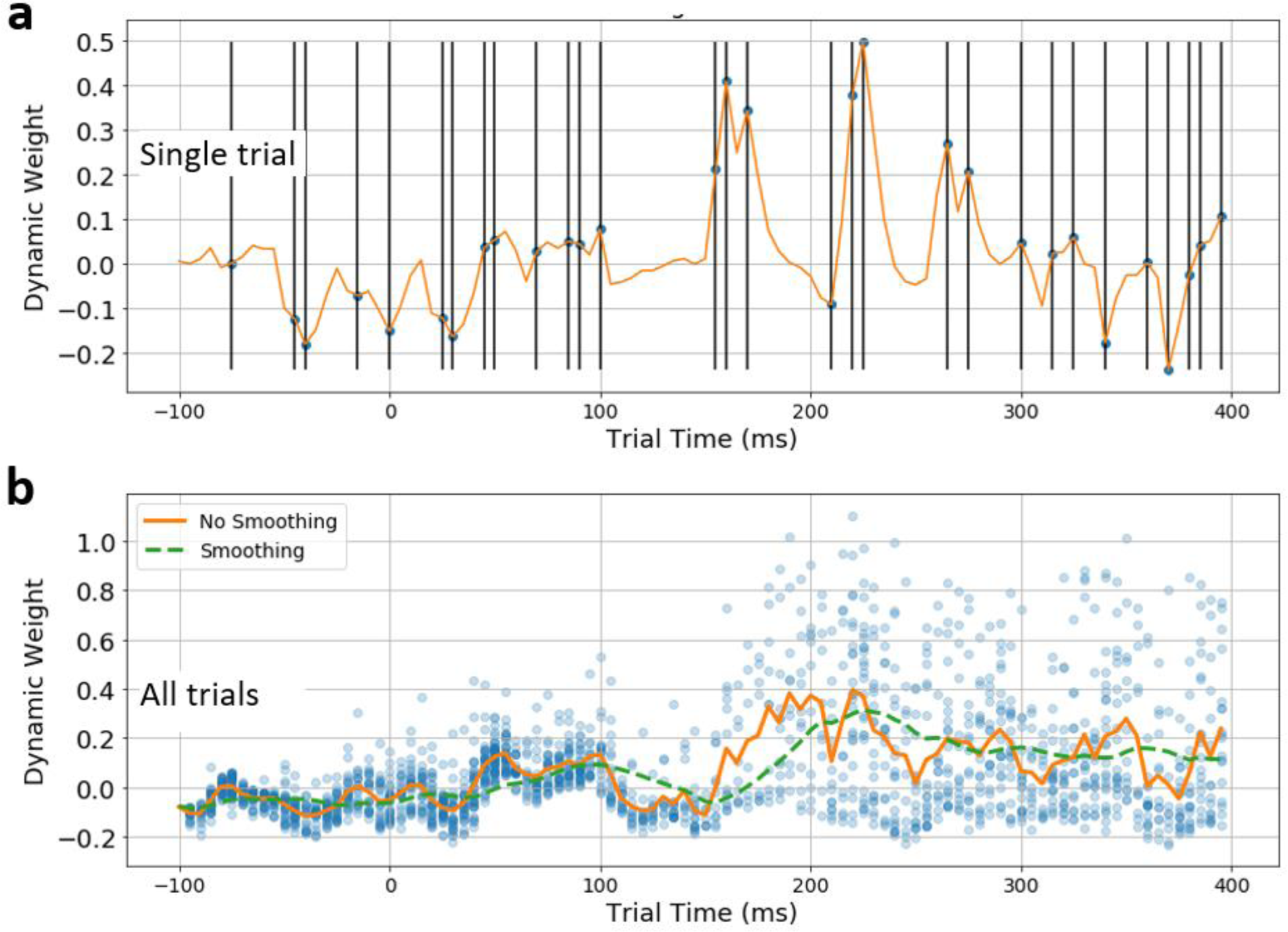
Selection of dynamic weight values and weights averaging. **A) Selection of weights which contribute to averaging**. Single trial spiking times for a given unit (black bars). Inferred dynamic weight values (orange curve) are selected at times when associated unit spikes (black dots), as the remaining weights do not contribute to the predicted spiking probability and therefore are not supervised to represent a meaningful value. **B) Averaging of weights**. Each blue point corresponds to a dynamic weight from a single trial. The orange curve represents a trial-average of every weight time-aligned with spikes. Green curve represents the Gaussian smoothing with a kernel including the 10 most recent bins.

Additionally, to compensate for the sparsity of spiking, we average these aggregated dynamic weights over the 10 most recent bins (Fig.9B).

#### Jitter correction

Jittering method ^15,30,41–43^ is often used in analysis of pairwise spiking correlations to minimize the influence of a potential stimulus-locked joint variation in population spiking that often occurs due to common input from many shared presynaptic neurons. To lessen the impact of such false positive (FP) connections in DyNetCP results and to ease the comparison with classical jitter-corrected CCG, we train a second static model on a jitter-corrected version of the data generated using a pattern jitter algorithm^41^ with a jitter window of 25ms. When the dataset is trial-aligned (e.g. VCN dataset), we use the correction procedure^41^ which determines the probability of a spike being present in each bin for all possible random shuffles of spikes among trials. For data which is not trial-aligned (e.g. synthetic dataset of HH neurons^25^), we randomly permute the spike times within the interval [−5ms, +5ms] before every weight update. New spike resampling is performed during every epoch of training. To assess putative connectivity, the jitter-corrected static weights *w*’_*j*→*i*_ produced by a jitter-corrected model are subtracted from the weights *w*_*j*→*i*_ trained on the original data to obtain 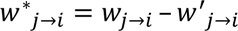 and 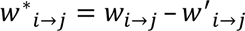.

### 2. Datasets

#### Synthetic network of HH neurons

We use a synthetic dataset consisting of Hodgkin-Huxley (HH) 800 excitatory and 200 inhibitory neurons with known ground truth (GT) synaptic connections^25^. The excitatory neurons influence 12.5% and the inhibitory neurons influence 25% of other neurons. We use the first 4320s of the 5400s-long data for training and the remaining 1080s for validation. Since DyNetCP requires training with trial-aligned data, we split each of these datasets into 1s “trials” consisting of 1 bins each. For training and evaluation we use spikes from the same subset of 20 excitatory neurons as was used for GLMCC validation^25^ (Fig.2.C,D,E).

#### Synthetic dataset with FP connections

Synchronous spiking at short time lags does not necessarily indicate a direct connection but can also be induced by a shared common input (Fig.10A) or an indirect serial chain connection (Fig.10B) among others. While it is notoriously difficult to distinguish these confounding correlations using descriptive statistical models^24^, the DyNetCP being a probabilistic model might lessen their impact by abstracting out such co-variates that do not contribute significantly to a prediction. To test the ability of DyNetCP to identify such confounds, we generate a synthetic dataset with known GT static and dynamic connectivity. The dataset consists of 100 excitatory neurons with their spikes generated by independent sampling from a Bernoulli distribution with spike rate between 0.002 and 0.01 per bin. The GT connection is active when spikes in neuron *n*_1_ at time *t* are followed by spikes in neuron *n*_2_ at a time *t* + *d* with a probability of 0.6. Randomly generated 1 “triplets” emulating common input and serial chain connections are added to the pool. he “triplets” are active either during the first and last 1 5 bin period or only during the middle 5 bin period. DyNetCP trained on this dataset (400 train and 200 validation trials, each 500 bins long) enables comparison of static and dynamic weights (Fig.10) that indicate that when the GT connection is present the **W**_**dyn**_ tends to be larger than **W**_**st**_, which enables classification of connections as TP and FP for both common input (Fig.10A) and serial chain (Fig.10B) connections for this synthetic dataset.

**Fig. 10.**
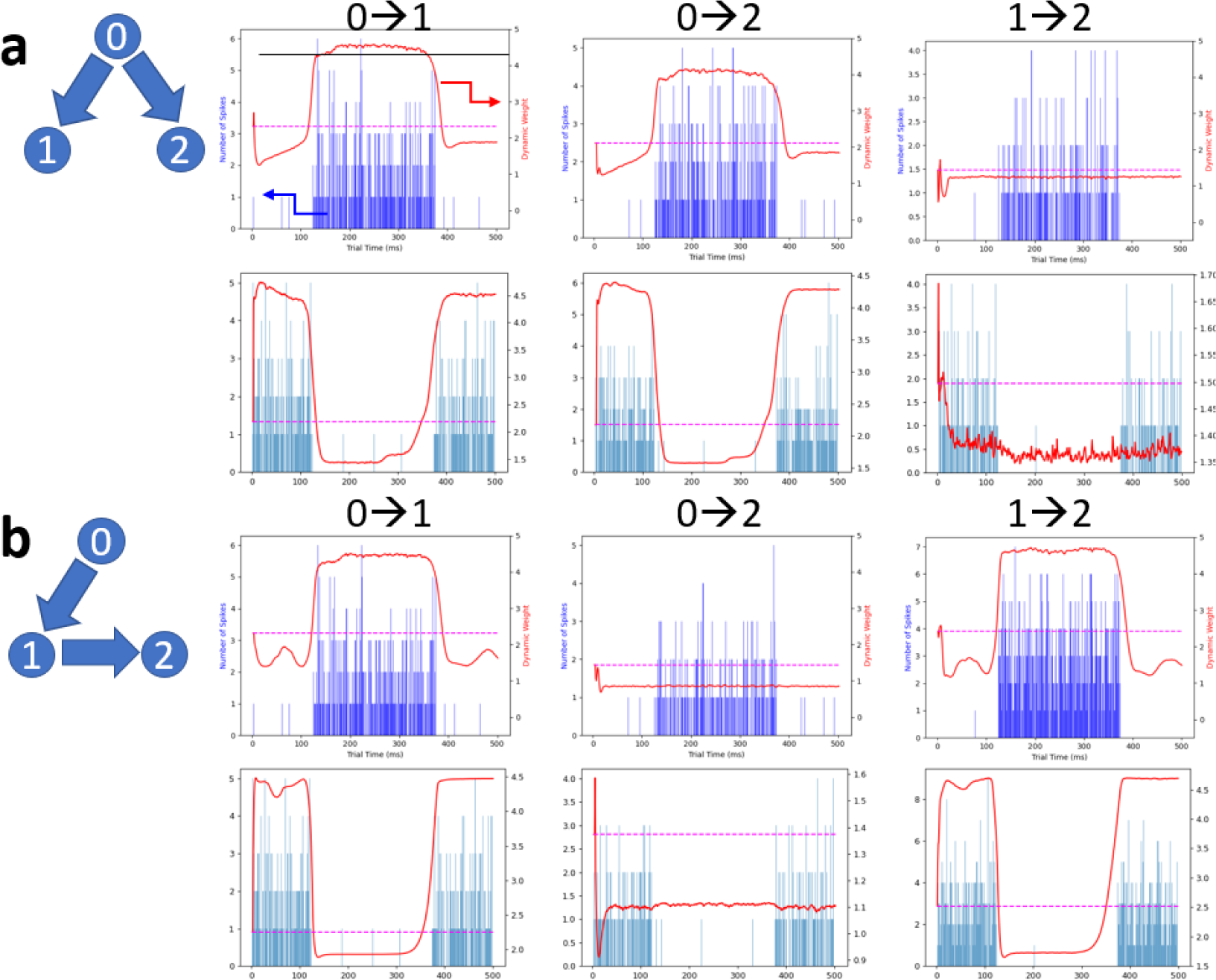
Performance of the model on synthetic dataset with FP connections. **A) Identification of FP connections for a “common input” triplet**. **Inset:** schematics of a triplet with common input from the unit 0 to both units 1 and 2. **Top row:** connection is active during the first and last 125 bin period of the trial. **Bottom row:** Connection is active only during the middle 375 bin period. Blue bars (left y-axis) represent a histogram of joint spiking probability (JPSTH anti-diagonal). Red curves correspond to inferred dynamic weights (right y-axis), while dotted magenta line represents static weight value. Comparison of dynamic and static weights might help to distinguish TP connections 0→1 and 0→2 from an artifactual FP connection 1→2. **B) Identification of FP connections for a “serial input” triplet**. **Inset:** schematics of a triplet with serial input from the unit 0 to 1 followed by 1 to 2. **Top row:** connection is active during the first and last 125 bin period of the trial. **Bottom row:** Blue bars (left y-axis) represent a histogram of joint spiking probability (JPSTH anti-diagonal). Red curves correspond to inferred dynamic weights (right y-axis), while dotted magenta line represents static weight value. Comparison of dynamic and static weights may help to distinguish TP connections 0→1 and 1→2 from an artifactual FP connection 0→2.

#### Experimental dataset recorded in visual cortex

To demonstrate DyNetCP performance with in-vivo high-density electrophysiological recordings, we used the publicly available Allen Brain Observatory-Visual Coding Neuropixels (VCN) database^15^ that contains spiking activity collected across 99,180 units from 6 different visual cortical areas in 58 awake mice viewing diverse visual stimuli. We use a “functional connectivity” subset of these recordings corresponding to the “drifting grating 5 repeats” setting recorded for animals (26 sessions in Table 1). Full-field drifting grating stimuli was passively viewed by a head-fixed animal freely running on a treadmill^15^. Stimuli consists of 2s presentation of drifting diffraction grating (temporal frequency 2 cycles/s, spatial frequency = 0.04 cycles per degree) that is interleaved with a 1s grey period (top left inset in Fig.1A). Presentation of gratings were repeated 75 times per orientation (0, 45, 90, and 135 degrees) and per contrast (0.8 and 0.4). To maximize the number of spikes available for training we use all angles and all contrast trials.

We split the 600 total experimental trials for each session into 480 training and 120 validation trials by selecting 60 training trials from each configuration of contrast or grating angle and 15 for validation. To further investigate the influence of brain state on connectivity dynamics we identify in each session trials with the average running speed above 5cm/s that are mar ed as “running”. he rest of the trials are mar ed as “not running”. When trained and validated, the model results can then be analyzed and compared for “running” vs “non-running” conditions separately.

### Models used

#### GLMCC model

Python code for GLMCC method is taken from the author’s version^51^. For these experiments, we use the synthetic dataset of interconnected HH neurons^25^. We focus specifically on recovering ground-truth (*GT*) excitatory connections. To facilitate a comparison with GLMCC model^25^, we here choose thresholds α_*w*_, α_*s*_, and α_*p*_ to maximize the Matthews Correlation Coefficient (MCC, Eq.9). MCC optimization balances between maximizing the number of *TP* pairs recovered while minimizing the number of *FP* pairs which are predicted.

Since a predicted connection is counted as a *TP* only once (i.e., *i* → *j* and j→ *i* do not count as two negatives if *i* is not connected to *j*) and the number of *GT* connections decreases when α_*p*_ increases, the MCC exhibits an increase beyond α_*p*_=100. A more detailed analysis of TP and FP rates as a function of pair spike threshold is presented in Fig.2 - figure supplement 1.

Though GLMCC performs well in recovering static network structure, we emphasize that GLMCC is not compatible with modeling network dynamics and thus cannot be integrated into DyNetCP. Specifically, our goal is to model the correlations of spiking of a network of neurons and track the changes of these correlations across time. GLMCC learns correlations as a function of the time lag relative to the spiking time of a reference neuron. It is hence unable to link correlations to specific moments in a trial time. Therefore, it cannot represent the dynamics of correlations, preventing its use in analysis of network activity relative to subject behavior.

#### Conditional Joint Peristimulus Time Histogram (cJPSTH)

The Joint Peristimulus Time Histogram (JPSTH) is a classical tool^39^ used to analyze the joint correlations of spiking behavior between two neurons and their variation across time. For a proper comparison with conditional DyNetCP results, we use a conditional joint peristimulus time histogram (cJPSTH), where we divide the columns of the JPSTH by the PSTH of neuron *i* (Fig.4B,C). Due to the sparsity of spiking in the data we use, it is beneficial to average the entries of the cJPSTH over short windows in time to more clearly extract the dynamics of the correlations between neurons. Specifically, we replace each entry (*a*, *b*) in the cJPSTH by the average of the *T* most recent bins having the same delay *a* − *b*, that is,

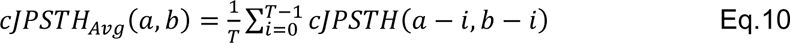

For most of the comparisons we use *T*=10 bins. Note that for most cJPSTH plots (e.g., Fig.4E), to directly compare to DyNetCP plots (e.g., Fig.4D), we only show data representing the “upper diagonal” of the full plot which represents data where neuron *i* spikes at the same time or before neuron *j* spikes. Specific implementation and comparisons of both models can be found in the accompanying Jupyter Notebook.

#### Dimensionality reduction

Dimensionality reduction^18,43,48^ is based on calculating principal components (PC) on the dataset and estimating the data variance that can be explained by a minimal number of PC. The dimensionality reduction code following a standard PCA reduction method^48^ is rewritten in Python and is used for analysis in Fig. 8A,B.

### 4. Statistical analysis of filtering and classification of units and pairs

The total number of units (raw units) recorded from visual cortices in 26 sessions of the VCN dataset^15^ is 18168. Units and pairs are filtered and classified by the following procedure.

#### Unit quality metrics

First, we apply standard unit quality metrics^15^ including amplitude cutoff < 0.1; violations of inter-spike interval (ISI) < 0.1; presence ratio > 0.95; isolation distance > 30; and signal to noise ratio (snr) > 1. This results in 5067 units (clean units).

#### Static weight peak threshold

Second, short-latency peaks within the first 10ms in the *w*^∗^_j→i_ and *w*^∗^_*i*→*j*_ of static weights are examined (example analysis for session 771160300 is shown Fig.2A,B). Following the accepted approach^15,19,21,25,30^ a putative connection is considered statistically significant when the magnitude of the peak exceeds a certain peak weight threshold α_w_ measured in units of standard deviations (SD) of the noise in the flanks, calculated within [51ms, 100ms] from both directions *w*^∗^_*i*→*j*_ and *w*^∗^_*i*→*j*_. When pairs are filtered with increasing peak weight threshold α_*w*_ (Fig.11A), the number of corresponding valid units is fast decreasing (black curve). For all 26 sessions 5067 clean units participate in formation of 5203 pairs with noticeable weight peaks. Filtering them using α_w_=7SD results in 1680 pairs corresponding to 1102 units. Close examination of these pairs (example analysis for session 771160300 is shown in Fig.11A) reveals that units not only form pairs (violet curve), but also form triplets (green), quintuplets (blue), and even participate in connections above 10-tuples (red). High radix hubs of this type are most likely FP artefacts imposed by sparse spiking (see below) and need to be filtered out. Indeed, electrophysiology spiking datasets are typically very sparse, with many interesting units often generating spikes at rates below 1Hz. Increasing the threshold α_*w*_ does help to filter out high radix connections, but still even for α_*w*_ as large as 8 there is a substantial number of units participating in connections larger than 3.

#### Unit spike rate

Therefore, as a third filtering step, we apply a unit spike rate threshold α_s_ to limit the minimal spike rate to 1Hz.

#### Pair spike rate threshold

It has been argued^25^ that, to reliably infer that a connection is present (i.e. to disprove the null hypothesis that a connection is absent), each binned time interval in the CCG histogram *d* should contain a sufficiently large number of spikes from both participating units. This implies that the spike rate of each neuron in a pair λ_*i*_ and λ_*j*_ should exceed the threshold α_*s*_, and the joint number of spikes in each time bin should exceed the pair spike threshold, defined as

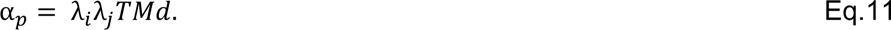

The confidence interval for a null hypothesis that two spike trains are independent stationary Poisson processes^25^ is then defined as

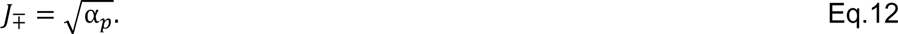

To disprove the null hypothesis and infer that a connection is statistically significant, α_*p*_ should exceed^25^ 10. To train the dynamic DyNetCP we use recordings binned at *d* of 5ms repeated over *M* of 480 randomized trials and the dynamic weights are influenced by *T*=10 most recent bins (Fig.9B). This corresponds to the *TMd* product of 24 that implies the minimum joint pair spike rate λ_*i*_λ_*j*_ of 0.41s^−2^. This threshold, however, might be insufficient for inference of the dynamic weight since only spikes within a short trial time interval are being used for learning. Indeed, analysis of an example pair (Fig.12A) that passed the α_w_=7SD threshold and has λ_*i*_λ_*j*_ = 0.73s^−2^, reveals that it exhibits on average just 2 spikes per trial or a total of 272 spikes for all 480 trials in a 500ms trial time window of interest. Corresponding jitter-corrected static weight matrix (Fig.12A, left) and cJPSTH (Fig.12A, bottom right) are very sparsely populated. The dynamic weight matrix (right top), although it is reproducing this sparseness, is not statistically convincing. Moreover, such pairs contribute significantly to high radix tuples.

By increasing the λ_*i*_λ_*j*_ pair spike rate threshold, the number of high radix tuples is decreasing (example analysis for session 771160300 is shown in Fig.11B), however the α_*p*_ threshold alone is not sufficient to eliminate them. Hence, for a given pair to be classified as a putative connection, the classification procedure requires the use of all three thresholds: α_*w*_, α_*s*_, and α_*p*_ (see Jupyter Notebook for specific implementation).

**Fig. 11.**
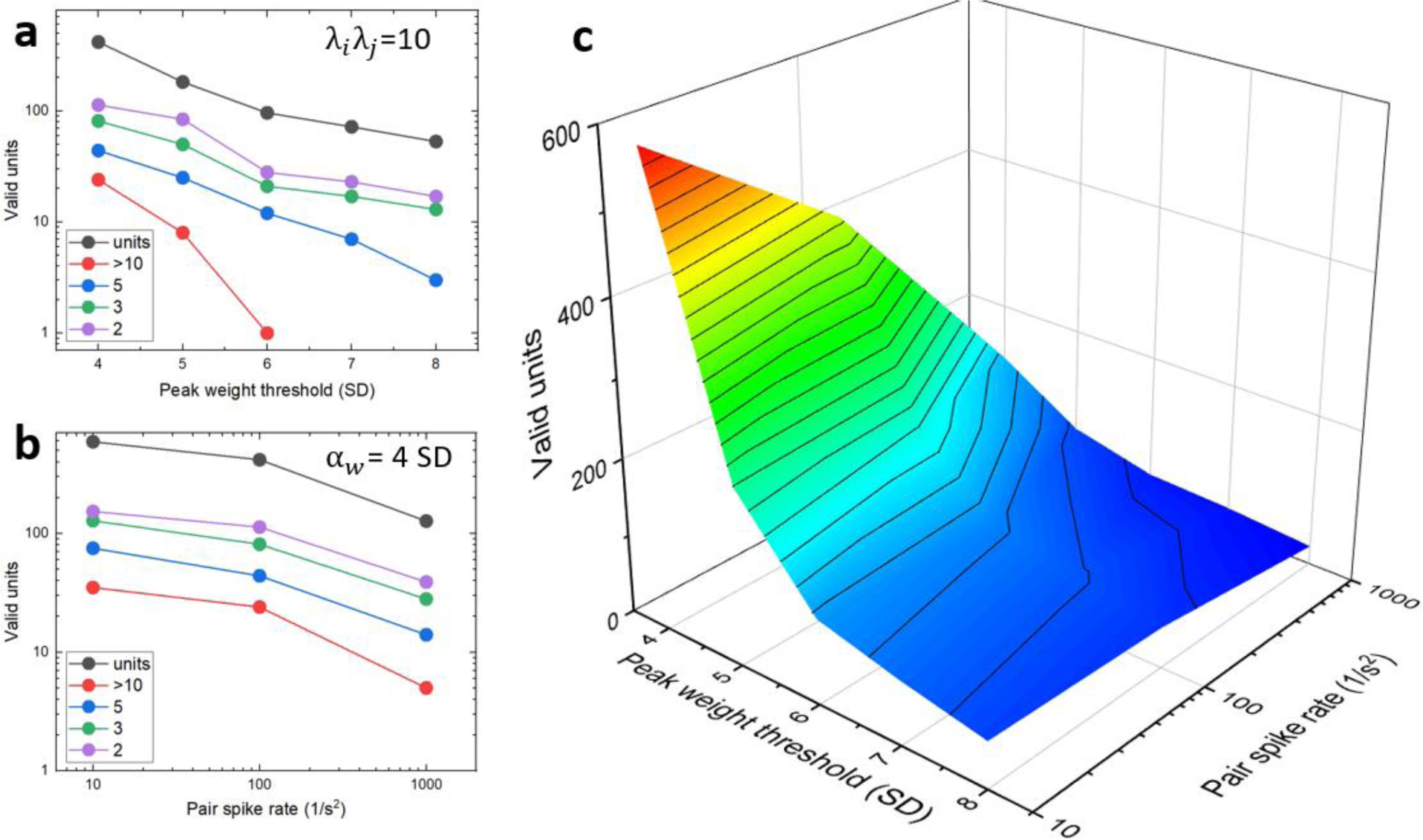
Thresholds for units and pairs filtering. **A)** Number of valid units (session #771160300) that form pairs with distinct static peak (black curve) for different peak weight thresholds. Violet, green, blue, and red curves correspond to units that participate in formation of pairs, triplets, quintuplets, and connections above 10-tuples, correspondingly. **B)** Number of valid units that form pairs with static peak above 4SD (black curve) for different pair spike rate. Violet, green, blue, and red curves correspond to units that participate in formation of pairs, triplets, quintuplets, and connections above 10-tuples, correspondingly. **C)** 3D plot of a number of valid units that form pairs for different peak thresholds and pair spike rates.

#### Choice of thresholds

When all three thresholds α_*s*_, α_*w*_ and α_*p*_ are applied simultaneously (Fig.11C) the number of valid units is decreasing rapidly with the increase of α_*w*_ to 7SD and λ_*i*_λ_*j*_ beyond 10 with slower decrease at higher thresholds. Examination of an example pair with joint pair spiking rate λ_*i*_λ_*j*_ of 50 s^−2^ (Fig.12B) indicates that while the static weight matrix shows a prominent (α_*w*_ >7SD) small latency peak, the cJPSTH is still populated relatively sparsely and the dynamic weight matrix resembles it much less (r_p_=0.27). In contrast, for an example pair with joint pair spiking rate λ_*i*_λ_*j*_ of 130 s^−2^ (Fig.12C) the resemblance between cJPSTH and the dynamic weight matrix is striking (r_p_=0.82, P_p_<0.001).

**Fig. 12.**
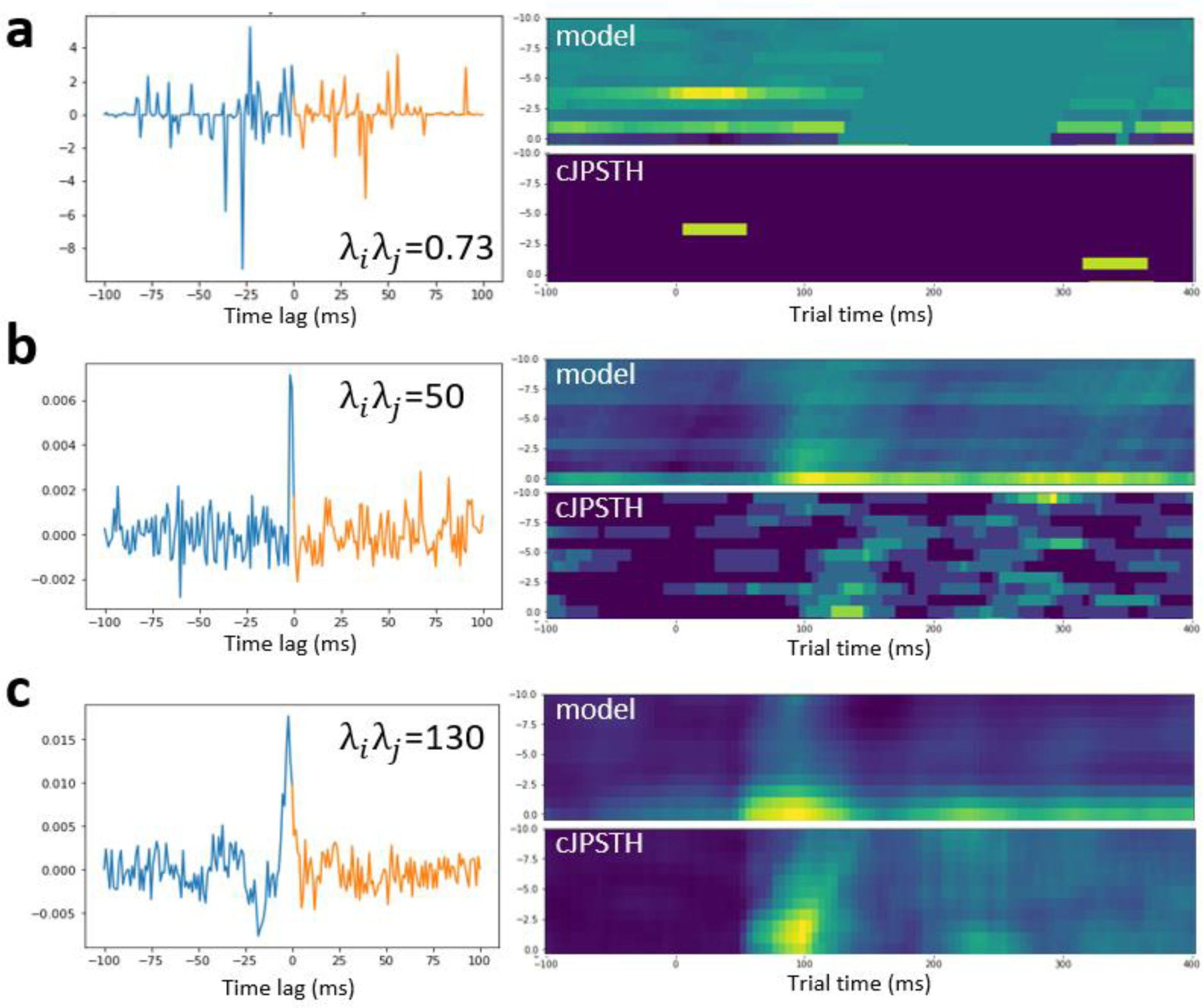
Example of pairs and dynamic model performance. **A)** Example of a pair of units (session #771160300, excitatory units from VISam and VISpm areas with spike rates 1.92Hz and 0.35Hz) that pass a peak weight threshold of 7SD (left static weight matrix plot), however, has pair spike rate as small as 0.73 s^−2^. Corresponding cJPSTH (bottom right) and dynamic weight plots (top right) are sparse. **B)** Example of a pair of units (session #771160300, excitatory VISp unit with 7.2Hz spike rate and inhibitory VISp unit with 6.8Hz rate) that pass a peak weight threshold of 7SD (left static weight plot), however, has moderate pair spike rate of 50 s^−2^. Corresponding cJPSTH and dynamic weight plots are shown on the right. **C)** Example of a pair of units that pass peak weight threshold of 7SD (left static weight plot), that has a relatively high pair spike rate of 130 s^−2^. Corresponding cJPSTH and dynamic weight plots are shown on the right.

For a fixed α_*s*_ of 1Hz and α_*w*_ of 7SD the distribution of all valid pairs with respect to their joint spike rates (Fig.13A) shows a maximum around λ_*i*_λ_*j*_=200 s^−2^ with a long tail to higher rates. To be conservative, for all the analysis of the inferred connectivity patterns in this work, we used λ_*i*_λ_*j*_ of 200 s^−2^ (α_*p*_=4800, red arrow in Fig.13A).

**Fig. 13.**
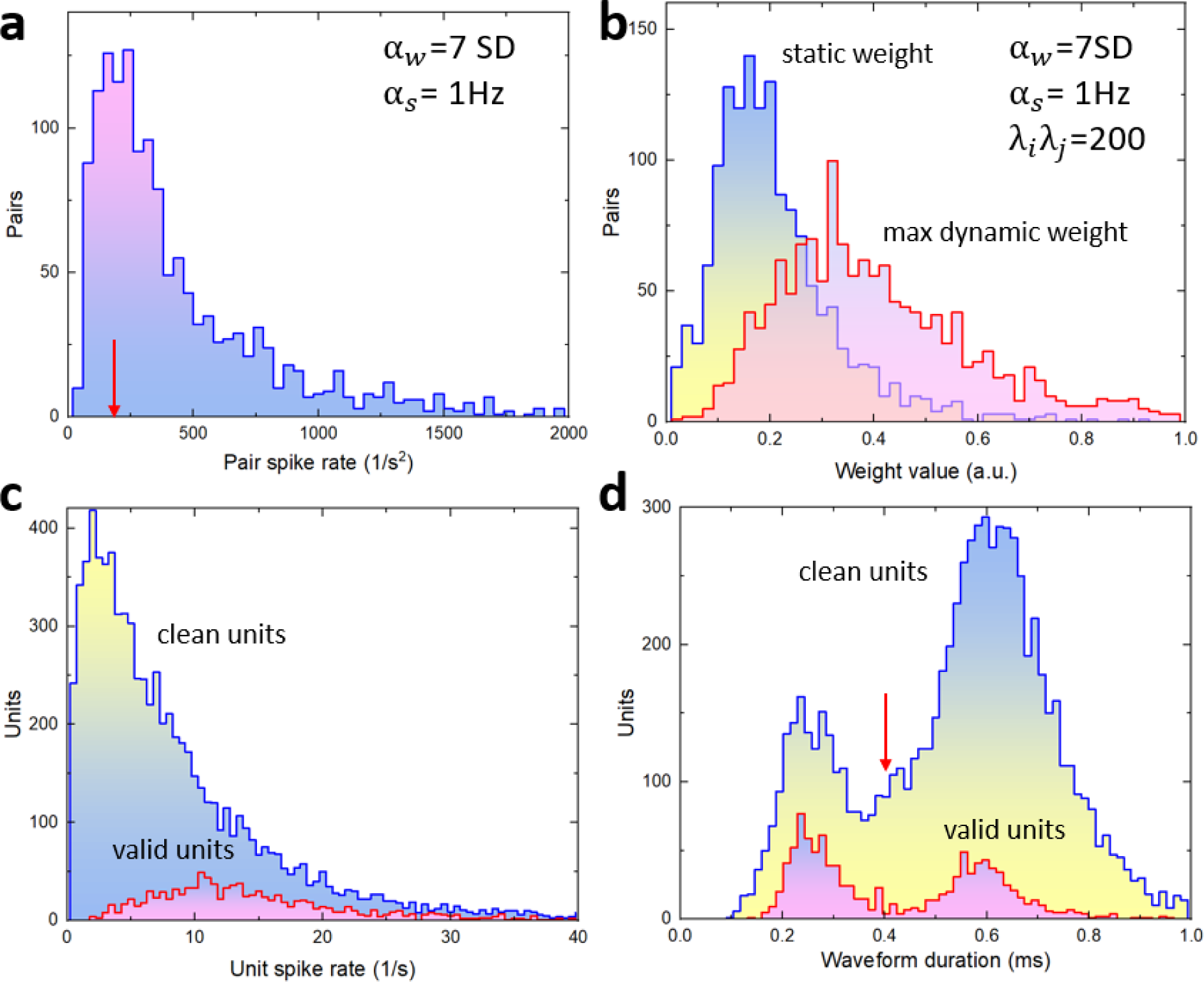
Filtered pairs and potential biases. **A)** Histogram of pairs recorded in all VIS cortices for all 26 animals that pass 7SD peak weight threshold and are formed from units with spike rate exceeding 1Hz. Red arrow shows the pair spike threshold of 200 s^−2^. **B)** Histograms of inferred static and maximum dynamic weights for filtered pairs. **C)** Histograms of unit spike rates for clean units and for valid units belonging to filtered pairs. **D)** Histograms of waveform duration for clean units and valid units. Red arrow indicates a threshold to classify units as inhibitory or excitatory.

Application of all 3 filters to all 26 sessions produces 808 units forming 1020 pairs (Table 1) with distribution of inferred static and dynamic weights shown in Fig.13B.

#### Units’ classification

Once all pairs are filtered, the directionality is assigned to each putative connection, either *j* → *i* or *i* → *j*, that is identified by the functional delay of the static weight peak binned at 1ms. Cortical depth perpendicular to cortical surface is calculated from unit coordinates in ccfv3 format (Allen Mouse Brain Common Coordinate Framework ^52^). Layer assignment is performed following method^15^ using units coordinates in the ccfv3 framewor and the annotation matri with reference space ey ‘annotation/ccf_ 1’. All units are classified as excitatory if their waveform duration is longer than 0.4ms and the rest as inhibitory (Fig.13D). Since transgenic mice were used for some of the sessions in the VCN dataset, optical activation of light-gated ion channel (in this case, ChR2) in Cre+ neurons with light pulses delivered to the cortical surface generate precisely time-aligned spikes that enable to link spike trains to genetically defined cell classes. Optical tagging was used to further classify inhibitory units^45^ as Parvalbumin-positive neurons (Pvalb), Somatostatin-positive neurons (Sst), and Vasoactive Intestinal Polypeptide neurons (Vip).

#### Potential biases

The statistical analysis and filtering procedure described above is biased towards strongly spiking units (Fig.13C) and as such can generate certain biases in connectivity patterns. For example, inhibitory units constitute about 18% of all 5067 clean units that pass unit quality metrics (Fig.13D). However, after filtering down to 808 valid units, they constitute over 50% of the population (Fig.13D). In general, inhibitory units have a higher spiking rate (for all clean units mean: 15.4Hz±0.29 SEM) than excitatory (mean: 6.3Hz±0.08 SEM). This might explain, at least partially, the over-representation of connections involving inhibitory units in the dynamic connectivity plots.

## Data Availability

All of the datasets used in this article are publicly available online under the Creative Commons Attribution 4.0 International license. Synthetic data of interconnected HH neurons^25^ is available at figshare^53^ (https://doi.org/10.6084/m9.figshare.9637904). The Allen Brain Observatory-Visual Coding Neuropixels (VCN) database^15^ is available for download in Neurodata Without Borders (NWB) format via the AllenSDK. The NWB files are available as an AWS public dataset (https://registry.opendata.aws/allen-brain-observatory/). Synthetic dataset with artifactual FP connections as well as trained static and dynamic weights for all 26 sessions of VCN database are availabel at Figshare https://doi.org/10.6084/m9.figshare.24879036

## Code Availability

DyNetCP introduced and used in this article is open-source software (GNU General Public License v3.0). The source code is available in a GitHub repository (https://github.com/NeuroTechnologies/DyNetCP) that contains codes for model training as well as a Jupyter notebook for generating the results used in the paper and in the SM.

## Notes

https://github.com/NeuroTechnologies/DyNetCP

### Competing Interest Statement

The authors have declared no competing interest.

### Summary of Updates

Text revised in response to peer review. Videos that were previously in Supplemental Materials, are included into the inline text.

